# Thymic selection of the T cell receptor repertoire is biased toward autoimmunity in females

**DOI:** 10.1101/2025.01.21.633929

**Authors:** Hélène Vantomme, Valentin Quiniou, Leslie Adda, Charline Jouannet, Vanessa Mhanna, Céline Albalaa, Pierre Barennes, Nicolas Coatnoan, Vimala Diderot, Johanna Dubois, Gwladys Fourcade, Kenz Le Gouge, Otriv Frédéric Nguekap Tchoumba, Martin Pezous, Paul Stys, Adrien Six, Encarnita Mariotti-Ferrandiz, David Klatzmann

## Abstract

Women represent about 80% of patients with autoimmune diseases. This may partly result from sex-based differences in T cell receptor (TCR) selection during thymocyte development, potentially influenced by hormones and the lower expression of the Autoimmune Regulator (AIRE) transcription factor in females.

To investigate this, we analyzed sex-specific differences in TCR generation and selection. We examined TCR repertoires in double-positive thymocytes and single-positive thymic cells, including CD8⁺ and CD4⁺ effector T cells and regulatory T cells (Tregs), derived from male and female organ donors. Minimal sex-based differences were observed in V and J gene usage, and there were no notable differences in TCR repertoire diversity, complementarity-determining region 3 (CDR3) length, amino acid composition, or network structure. No TCR sequences were exclusive to either sex.

However, female effector T cells exhibited a significantly higher prevalence of TCRs specific to self-antigens implicated in autoimmunity compared to males, while female Tregs showed a reduced frequency of such TCRs. These differences were not observed for TCRs targeting self-antigens unrelated to autoimmunity or antigens associated with cancer or viruses.

Our findings identify a sex-specific imbalance in thymic selection of TCRs with autoimmunity-associated specificities, providing mechanistic insight into the increased susceptibility of women to autoimmune diseases.

## Introduction

Several studies have highlighted a sex imbalance in diseases involving the immune system. This is particularly evident in the case of autoimmune diseases (AIDs), where 80% of affected individuals are women, and the severity of the condition varies between the sexes (1–3). In addition, men are more severely affected by infectious diseases such as COVID-19 and tuberculosis (4,5). The response to preventive immunotherapies such as vaccination, and to curative treatments, like immune checkpoint inhibitors or anti-TNF alpha, is also sex dependent (6). These observations indicate the presence of intrinsic biological differences in immune responses between males and females.

A variety of factors have been identified that could help explain the observed sex-based differences. Among these, sex hormones have been shown to influence immune responses by acting on immune cells that express specific receptors to these hormones (7–9). Sex hormones could directly influence the adaptive immune response at the level of thymocyte differentiation and selection by acting on the expression of Autoimmune Regulator (AIRE) in thymic epithelial cells. AIRE is responsible for the thymic expression of otherwise tissue-specific antigens, contributing to both the negative selection of effector cells and the positive selection of Tregs that recognize such antigens (10). Research has shown that testosterone increases AIRE expression in medullary thymic epithelial cells, whereas estradiol decreases it (11,12). Furthermore, a recent study has shown that the transcriptomic profiles of these specialized antigen-presenting cells in the thymus differ between males and females (13). This suggests that there may be differences in the selection of the T cell receptor (TCR) repertoire between males and females.

Despite these findings, there is still limited knowledge about potential differences in the TCR repertoire between the sexes. The generation of the TCR is a key process in T-cell development that takes place in the thymus. The TCR consists of two chains, alpha (TRA) and beta (TRB), each of which resulting from a random genetic rearrangement involving the V (variable), D (diversity) [for the TRB], and J (joining) genes. The combination of these gene segments is accompanied by insertions and deletions, creating a highly diverse region, the Complementarity Determining Region 3 (CDR3) (14). It is this region of the TCR that predominantly interacts with the peptide, a critical interaction for antigen recognition and subsequent T cell activation. The ability to specifically recognize diverse antigens via the CDR3 is central to the efficiency and specificity of the adaptive immune response.

TCRs are randomly generated, which means that while some may be useful, others may be harmful by recognizing self-antigens and causing autoimmunity. The current understanding is that during thymocyte development, selection is based on the avidity of their TCRs for antigens presented by cortical and medullary thymic epithelial cells. Thymocytes expressing TCRs with insufficient avidity fail to receive survival signals and undergo death by neglect during positive selection, whereas those with excessive avidity are actively eliminated via negative selection to ensure self-tolerance. In contrast, thymocytes with intermediate avidity are positively selected and mature into single-positive cells, either CD4+ or CD8+. Thymocytes with the highest of these intermediate affinities develop in Tregs. Given AIRE’s pivotal function in these processes, sex differences in AIRE expression may significantly influence thymic selection.

The TCR repertoire of female and male peripheral T cells have shown that their diversity, particularly that of the TRB repertoire, is influenced not only by the age of the individuals but also by their sex (15–18). However, another study showed no association between the sex of individuals and the diversity of their TCR repertoire (19). Nevertheless, the study of peripheral blood TCRs reflects that of an immune system that has undergone multiple immune responses. For example, observed differences between males and females in their responses to self-antigens associated with autoimmunity could indicate either the cause (the TCRs drive the disease) or the consequence (the TCRs are expanded because of the disease).

To assess whether there are significant differences in the generation and thymic selection of the TCR repertoire between males and females, we therefore focused on analyzing the TCR repertoires of thymocytes. These cells represent the latest stage in T cell development and have not yet been involved in immune responses. Their repertoires thus represent the mechanisms of TCR generation for the analysis of CD4/CD8 double positive (DP) cells, and of TCR selection for CD4 and CD8 simple positive (SP) T cells. We generated a unique data set comprising the sequences of the TRA and TRB chains of DP cells, CD8 and CD4 SP effector cells, and CD4 SP Tregs from organ donors, both infants and adults. The analysis of TRA and TRB repertoires using state-of-the-art strategies revealed no sex differences in repertoire generation (DP cell repertoire), but an enrichment in females for CD8 SP T cells bearing CDR3 associated with specificities related to autoimmunity.

## Results

We analyzed thymic TCRαβ repertoires from 22 human organ donors (11 females, 11 males, aged between 73 days and 64 years) (**Figure 1, Supplementary Figure 1**). For each donor, CD3+CD4+CD8+ double positive (DP), CD3+CD4-CD8+ CD8 single positive (SP), and CD3+CD4+CD8-CD4 SP thymocytes were sorted by flow cytometry. In 16 donors, CD4 SP cells were further subdivided into CD25-conventional CD4 Teff and CD25+ CD4 Treg subsets, whereas in the remaining six donors, total CD4 SP cells were analyzed as Teff-enriched populations. This resulted in 20 DP, 21 CD8 SP, 22 CD4 Teff SP and 14 CD4 Treg SP samples (**Figure 1, Supplementary Figure 1**). Bulk TCR sequencing was then performed on each cell subset, with the aim to characterizing the TRA and TRB repertoires. Rank–frequency plots of clonotype abundances displayed comparable heavy-tailed distributions across donors, without any obvious systematic differences between males and females (**Supplementary Figure 2**).

**Figure 1:**
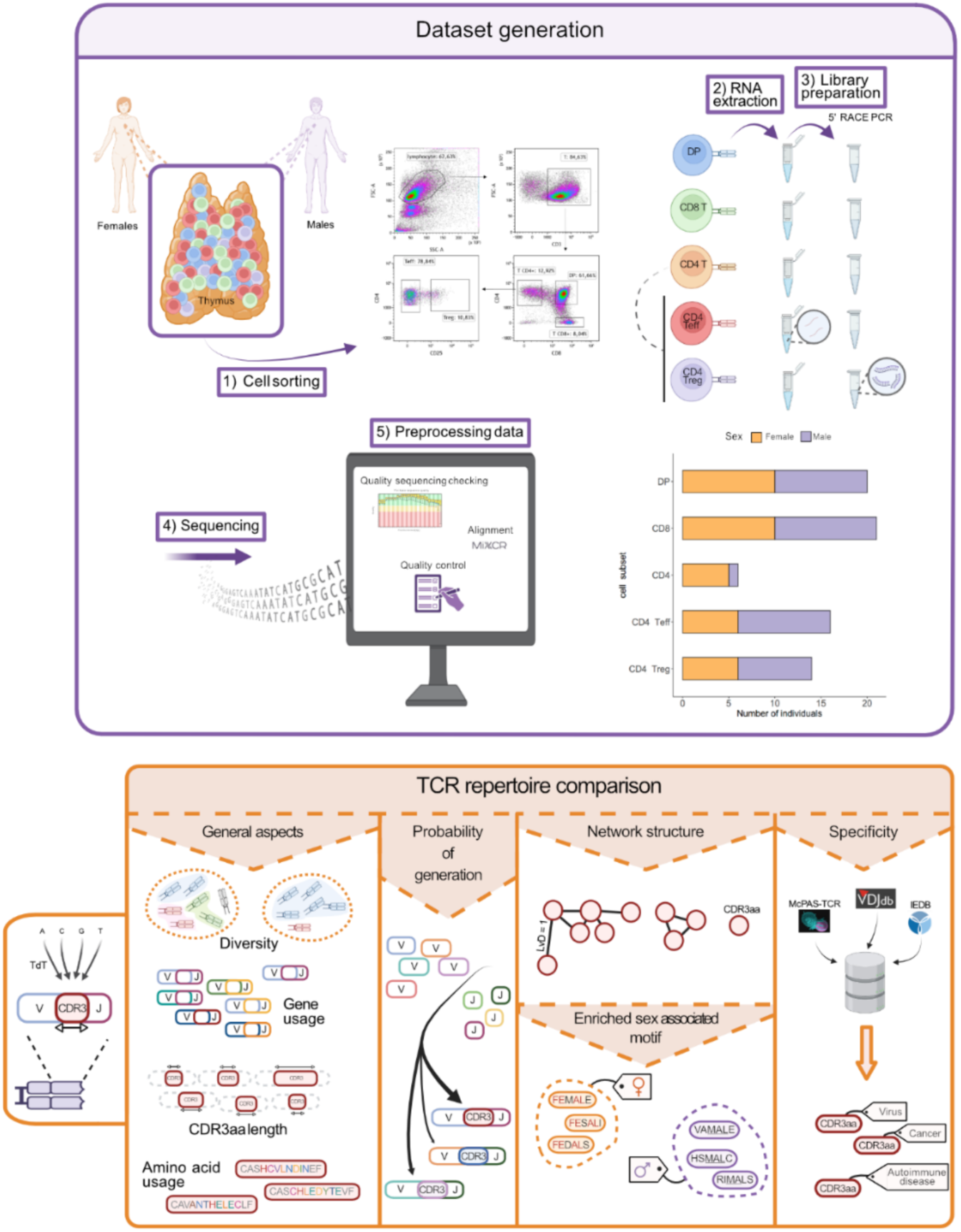
Schematic overview of the generation of the thymic TCR dataset and the analytical pipeline. Partially created with Biorender.com **Top Panel:** Generation of the thymic TCR dataset. From deceased human thymuses of males and females, we isolated key T-cell subtypes through cell sorting. These subtypes included double-positive (DP) cells (CD3+ CD4+ CD8+), single-positive (SP) CD8+ cells (CD3+ CD4-CD8+), SP CD4+ cells (CD3+ CD4+ CD8-) that were further separated into T effector (Teff, CD3+ CD4+ CD8-CD25-) and Treg (CD3+ CD4+ CD8-CD25+) cells. TCR libraries were generated from the RNA of each cell population using rapid amplification of cDNA ends by PCR (5’RACE PCR). Following sequencing, data preprocessing involved quality sequencing checks, contig alignment, and quality control. The final dataset comprised 20 DP samples (male-to-female ratio of 1:1), 21 SP CD8+ samples (1.1:1), 6 SP CD4+ samples (1:5), 16 SP CD4 Teff samples (1.67:1), and 14 SP CD4 Treg samples (1.33:1). Males are depicted in violet and females in orange. **Bottom Panel:** Analytical pipeline. We compared the TCR repertoires of males and females across various dimensions. We evaluated general aspects of the TCR repertoire were evaluated, including diversity, gene usage, CDR3aa length distribution, and aa usage within the CDR3 region. Additionally, we analyzed the probability of sequence generation and the TCR repertoire structure based on CDR3aa sequence similarity. We identified differentially expressed TRB CDR3aa motifs between sexes and analyzed TRB CDR3aa sequence specificity.

### Minimal sex-based differences in TCR V and J gene usage across thymic cell subsets

The first level of diversity in generating TCRs is the usage of V and J genes (20). We analyzed the usage of TRA and TRB V and J genes and their combinations in each thymocyte population (**Figure 1**).

Males and females had a relatively similar V gene usage, as evidenced by the lack of a clear separation in the Principal Component Analysis (PCA), for both TRA and TRB, and across all cell subtypes studied (**Figure 2A**). However, we observed some individual differences in V and J gene usage between males and females, with some being specific to certain cell subtypes (**Supplemental Figures 3-6**), and others observed across multiple cell subtypes (**Supplemental Figures 3-6**). For example, TRBV6-5 was found to be more highly expressed in females compared to males in DP cells (**Supplemental Figure 3C**) and CD8 SP cells (**Supplemental Figure 4C**).

**Figure 2:**
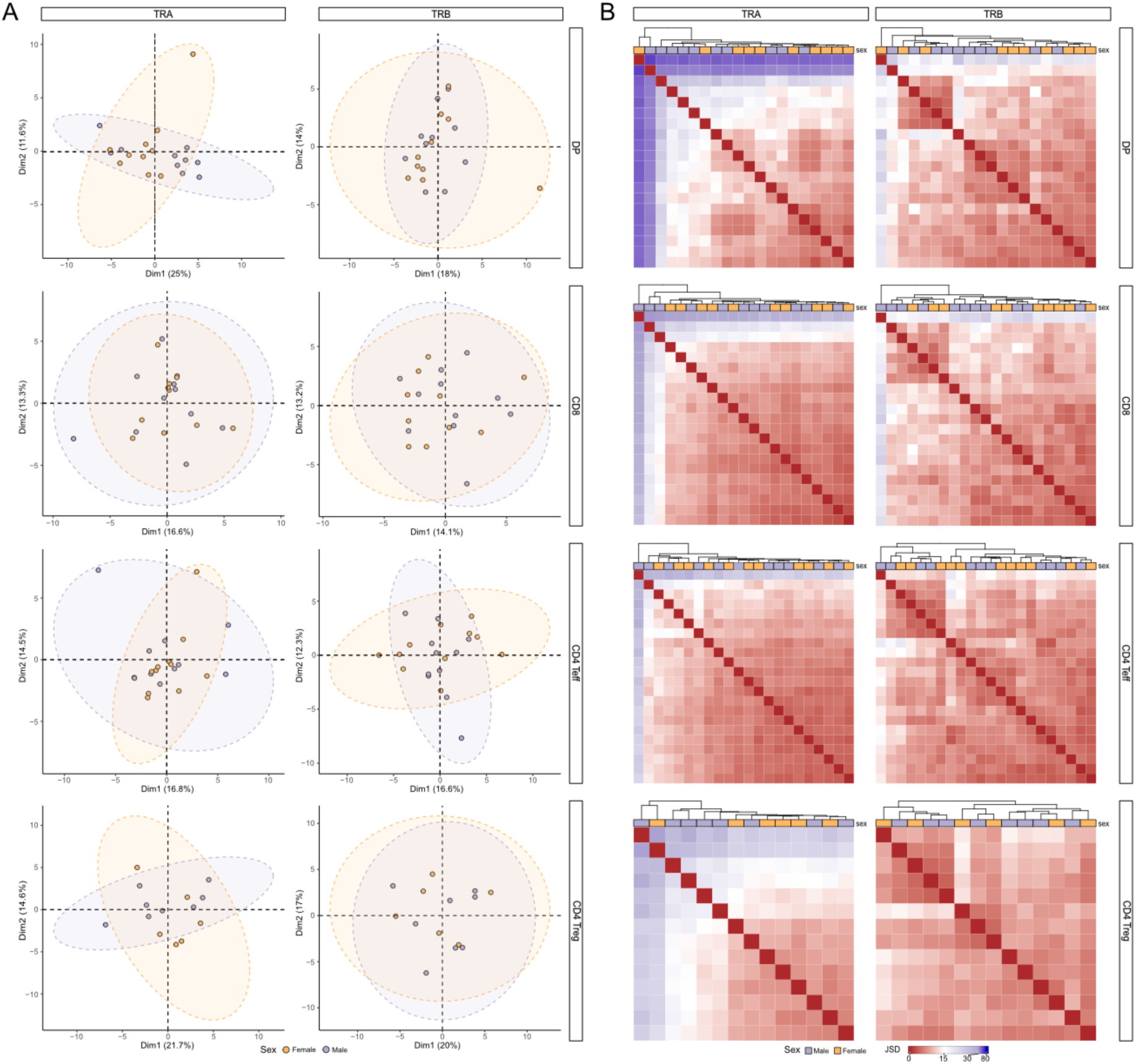
Comparable overall TCR gene usage between males and females. **(A)** Principal Component Analysis (PCA) derived from the distribution of TRAV (left) and TRBV (right) gene usage frequencies across sex groups (males vs females), showing results for DP (N = 22), CD8 (N = 23), CD4 Teff (N = 24) and CD4 Treg (N = 14) cells (displayed from top to bottom). Each point on the graph represents an individual. Ellipses indicate 95% confidence intervals. **(B)** Heatmap showing the Jensen-Shannon Divergence (JSD) score between samples, derived from the distributional usage of TRAV-TRAJ (left) and TRBV-TRBJ (right) gene associations in DP (N = 22), CD8 (N = 23), CD4 Teff (N = 24) and CD4 Treg (N = 14) cells (displayed from top to bottom). Hierarchical clustering was performed using the Euclidean distance and the complete linkage method. Males are shown in violet; females in orange.

In the following step, we compared the usage of TRAV-TRAJ and TRBV-TRBJ gene combinations across all individuals using the Jensen-Shannon Divergence (JSD) score (**Figure 2B**). The hierarchical clustering showed no clear separation between male and female individuals for any of the chain or cell subtype, indicating no significant differences in the gene combination usage between males and females (**Figure 2B**). A similar observation was made when directly analyzing the frequency of these V-J gene combinations (**Supplemental Figures 3E-6E, 3F-6F**).

### Comparable TCR repertoire diversity between males and females

A high diversity of the TCR repertoire is essential for generating a broad potential reactivity against a variety of antigens. We thus compared the TCR repertoire diversity between males and females. To achieve this objective, we analyzed Rényi diversity curves (from zero to infinite 𝛼 parameters) that measure various aspects of TCR repertoire diversity. At different 𝛼 values, the Rényi index places more or less weight on the contribution of the most frequent clonotypes. Higher 𝛼 values highlight the most prevalent clonotypes, thereby revealing the effect of expanded clones on overall diversity.

Rényi curves showed comparable diversities between sexes in DP and CD4 Teff cells, for both TRA and TRB, and a difference in CD8 and CD4 Treg cells (**Supplemental Figure 7**). In CD8 cells, the curves begin to diverge at low 𝛼 values (**Supplemental Figure 7**), with an earlier inflection in males, indicating a less balanced distribution of TCR clones. This suggests that in males, fewer clones dominate the repertoire at lower diversity thresholds compared to females. In Tregs, while the inflection point is similar, the difference seems to be characterized by higher diversity and richness in females compared to males (**Supplemental Figure 7**). However, no significant differences were observed for the Shannon diversity index (𝛼 = 1), the Simpson index (𝛼 = 2), and the Berger-Parker index (𝛼 = infinite), for both chains and all cell subtypes (**Figure 3**). These results indicate that the diversity of thymic TCR repertoire is similar between males and females.

**Figure 3:**
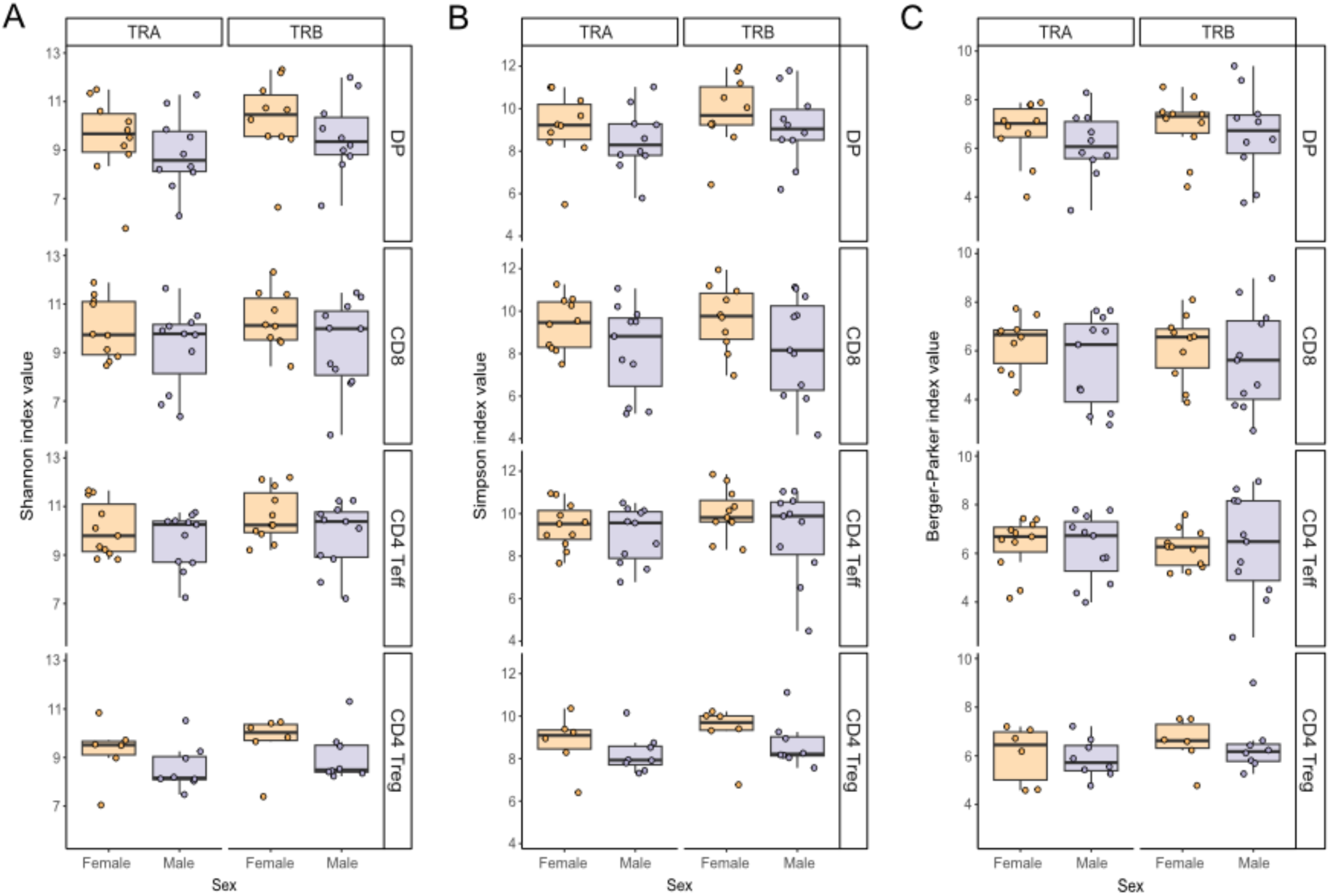
Comparable thymic TCR repertoire diversity between males and females. Boxplots display Shannon **(A)**, Simpson **(B)**, and Berger-Parker **(C)** index values for TRA (left) and TRB (right), across thymic T cell subtypes, displayed from top to bottom, in DP (N = 22), CD8 (N = 23), CD4 Teff (N = 24) and CD4 Treg (N = 14). Each point on the graph represents the median value from 50 rarefactions per sample. Statistical analysis (Wilcoxon test) showed no significant sex bias in TCR repertoire diversity (p > 0.05). Males are shown in violet; females in orange.

### Comparable CDR3aa length distribution in male and female TCR repertoires

Most of the repertoire diversity is generated by the random addition of nucleotides within the VDJ junctions, resulting in the hypervariable CDR3 region that interacts with the peptide-MHC complex, and is mostly responsible for the TCR specificity. The CDR3 length has been shown to directly influence the TCR’s ability to interact with the peptide (21). We thus studied the distribution of CDR3 aa (CDR3aa) sequence lengths between males and females to identify potential TCR generation and/or selection biases (**Figure 1**).

Our results showed that males and females have a comparable distribution of CDR3aa lengths for both TRA (**Supplemental Figure 8**) and TRB (**Figure 4A**). CDR3aa sequences in TRA were shorter than those in TRB, as previously described in peripheral blood T cells (22,23). Although there were subtle differences in certain CDR3aa lengths between males and females, these variations were minimal, involving sequences that represent less than 2% of the total TCR repertoire (**Figure 4A** and **Supplemental Figure 8**). Altogether, the overall distribution of CDR3aa length is comparable between males and females.

**Figure 4:**
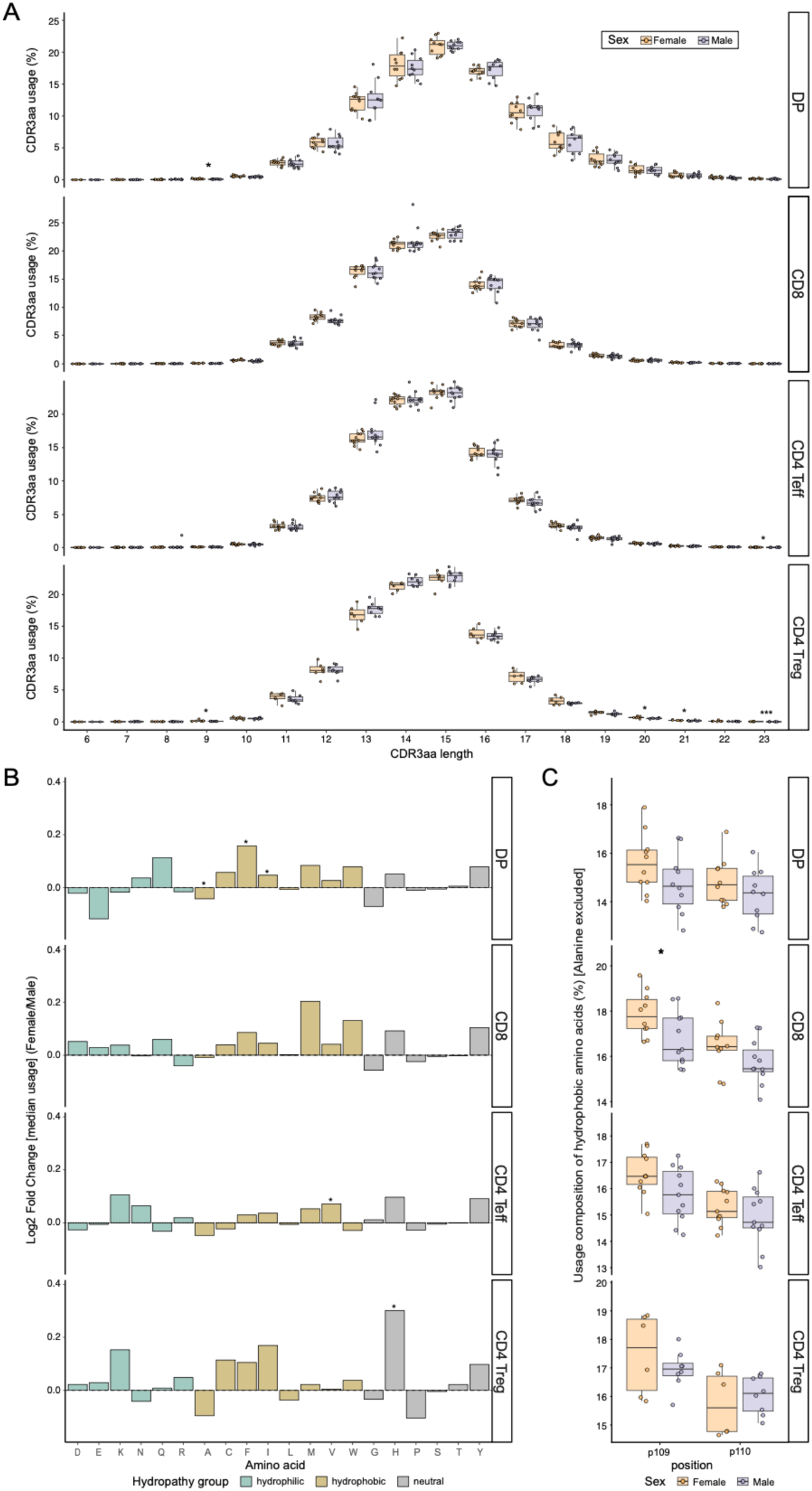
CDR3aa length and amino acid composition of TRB CDR3s in males and females. **(A)** Distribution of TRB CDR3aa length usage in DP (N = 22), CD8 (N = 23), CD4 Teff (N = 24) and Treg (N = 14) SP cells (displayed from top to bottom). Stars indicate statistical differences between males and females based on the p-value of the Wilcoxon test (*: p < 0.05, **: p < 0.01). **(B)** The data on amino acid (aa) usage between males and females is presented as the log2 fold change of the median per-donor usage in females over males for each aa in the p108 to p114 CDR3aa region for TRB. A line at log2 fold change = 0 is indicative of the direction of the difference in usage frequency. The bars are colour-coded to the hydropathy class of the aa as defined by the Kyte-Doolittle-based IMGT classification (neutral aa by gray, hydrophilic aa by blue-green and hydrophobic aa by gold). Stars indicate statistical differences of usage between males and females based on the p-value of the Wilcoxon test (*: p < 0.05, **: p < 0.01). **(C)** Position-specific usage of hydrophobic aa (excluding alanine, due to its weak hydrophobicity) at IMGT positions p109 and p110 in TRB across thymic T cell subtypes, including DP (N = 22), CD8 (N = 23), CD4 Teff (N = 24) and CD4 Treg (N = 14). For each donor, the values represent the proportion of unique TRB CDR3aa sequences carrying a hydrophobic amino acid at the indicated position. Stars indicate significant sex differences based on the p-value of the Wilcoxon test (*: p < 0.05), with a significant increase in hydrophobic usage at p109 in female CD8 SP cells. Males are depicted in violet; females in orange.

### Subtle differences in the composition of CDR3aa in the thymic TCR repertoires of males and females

We analyzed the aa usage within the FG loop of the CDR3, which is in contact with the peptide-MHC complex (24). The classification of aa was determined based on their hydropathy properties, categorized as follows: hydrophilic, hydrophobic, or neutral (25). We observed significant differences in the usage of some aa between males and females for both TRA (**Supplemental Figure 9**) and TRB (**Figure 4B**). In DP cells, these differences predominantly involved hydrophobic aa: females exhibited higher alanine (A) usage in TRA and lower in TRB compared with males (**Figure 4B** and **Supplemental Figure 9**). Additionally, females showed increased usage of phenylalanine (F) and isoleucine (I) in TRB (**Figure 4B**). Furthermore, other aa with varying hydropathy properties were differentially used between males and females in CD4 Teff and Treg populations (**Figure 4B** and **Supplemental Figure 9**).

We then examined the usage of aa specifically at positions p109 and p110 of the TRB CDR3aa region, as hydrophobic aa at these two positions have been associated with a stronger recognition of self-antigens (26–28). For each individual, we calculated the proportion of unique TRB CDR3aa carrying a hydrophobic aa (excluding alanine, which is only weakly hydrophobic) at these two positions. This positional analysis revealed a significant increase in the usage of hydrophobic amino acid at position p109 in female CD8 cells compared with males, with similar trends observed at p109 in DP and CD4 Teff subsets and at p110 in CD8 cells (**Figure 4C**). Taken together, these positional patterns suggest a subtle sex-biased enrichment of hydrophobic amino acids at key TRB CDR3 positions that could be compatible with a slightly increased potential for self-reactivity and cross-reactivity in female CD8 T cells (28).

### Slight sex-based variations in thymic TCR generation probability distribution

Another approach to evaluating biases in the generation and selection of the TCR repertoire is to compare the probability of generation (Pgen) of CDR3 nucleotide sequences (**Figure 1**). Pgen represents the likelihood of a sequence being produced, according to models of V(D)J recombination (14,29). For DP and CD8 cells only and for each TCR chain, a sequence generation model was created using 100,000 non-productive nucleotide sequences. Using this model, the Pgen of all productive TCR sequences that make up each repertoire was calculated.

When comparing the distribution of Pgen between males and females, we observe a very slight difference of distribution, characterized by a nearly null Kolmogorov-Smirnov score (D_K-S_ < 0.05) (**Figure 5**). This difference is more pronounced in the TRB compared to the TRA chain (**Figure 5**). As this minor discrepancy might be indicative of variations among individuals irrespective of their sex, we have generated 20,000 permuted mixed-sex groups utilizing our dataset. Our findings revealed a significant distribution difference of Pgen sequences between these mixed groups population (DP – TRA: D_K-S_ = 0.0183 ± 0.0145 (µ ± sd), DP – TRB: D_K-S_ = 0.0190 ± 0.0161, CD8 – TRA: D_K-S_ = 0.0227 ± 0.0153, and CD8 – TRB: D_K-S_ = 0.0259 ± 0.0140). However, the observed difference in the Pgen distribution between males and females is higher than that between the mixed groups in the DP TRB.

**Figure 5:**
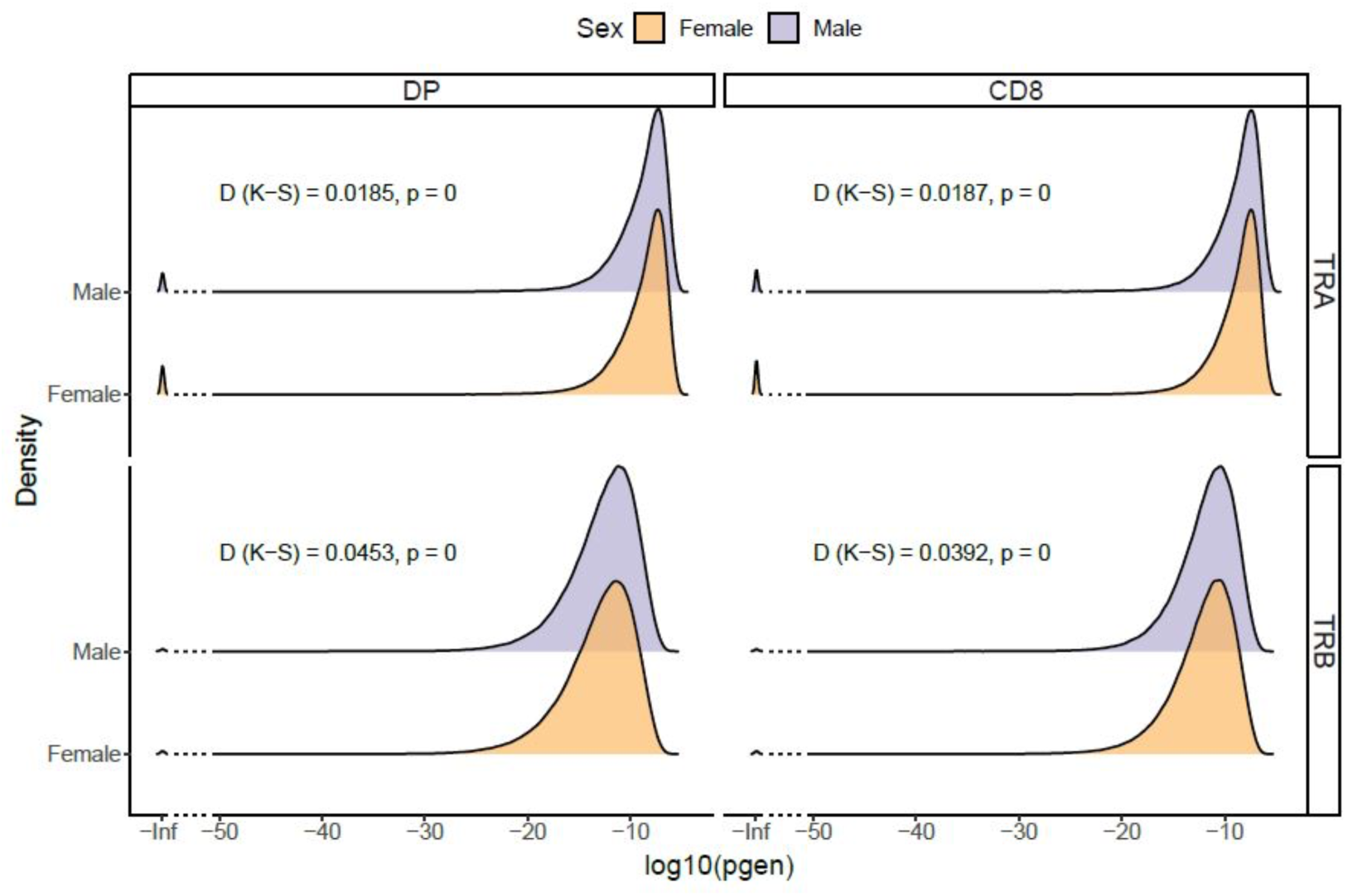
Probabilities of generation of TCRs in males and females. The figure shows the distribution of Pgen (probability of generation) sequences between males and females for TRA and TRB in DP and CD8 cells. A V(D)J recombination model was created using 400,000 non-productive random sequences derived from the nonproductive sequences of all individuals for each chain, both for DP and CD8 cells (65). This model was then used to calculate the Pgen of sequences for each individual (66). The overall distribution comparison between males and females was tested using the Kolmogorov-Smirnov test, with the D value and associated p-value indicated. Males are depicted in violet and females in orange.

**Figure 6:**
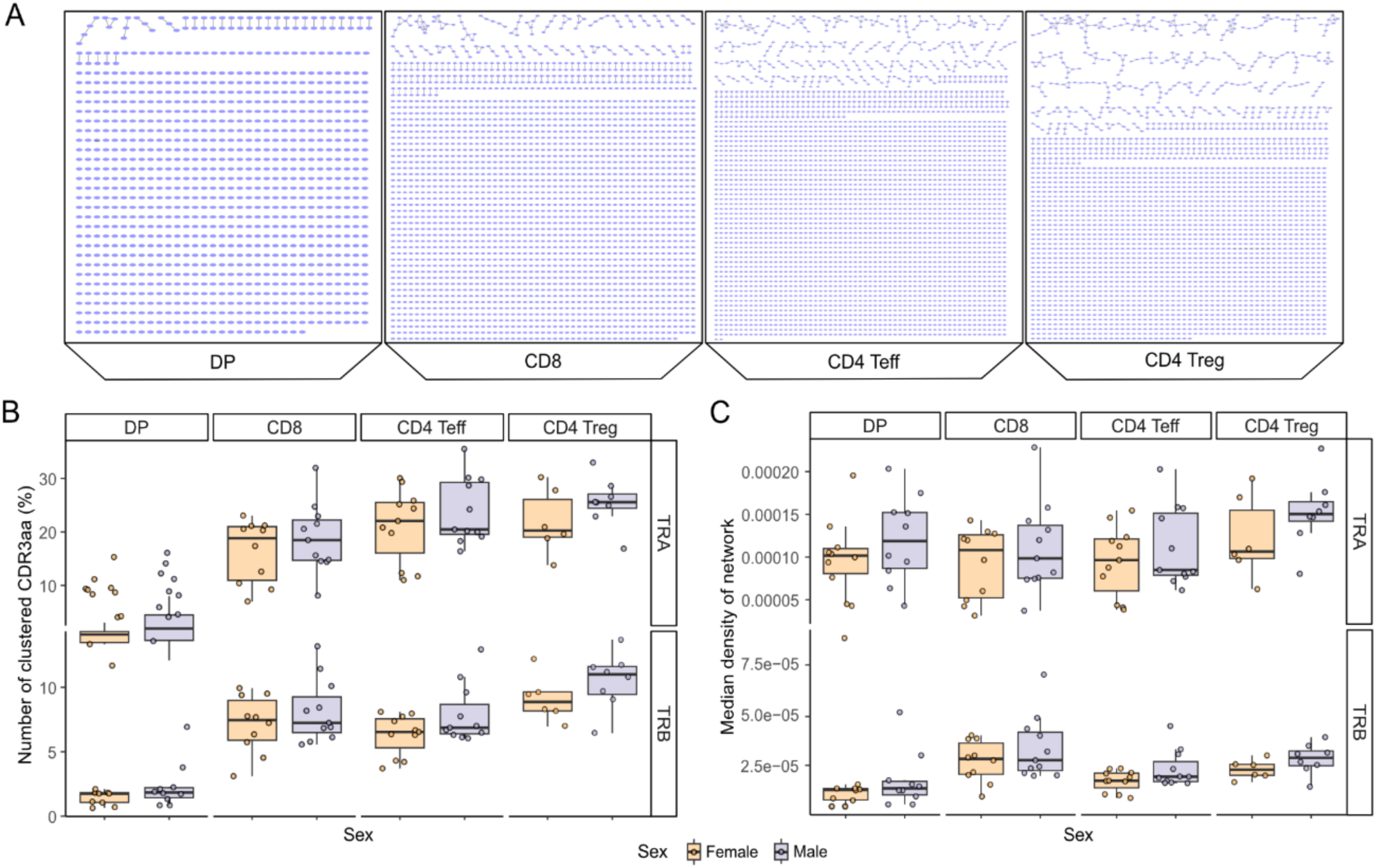
Comparable TCR repertoire network structure based on CDR3 amino acid sequence similarity between males and females. For each sample, 100 random subsamplings were performed on the minimum number of CDR3aa per cell subtype. Two CDR3aa are linked if they have a Levenshtein distance of one. **(A)** TRA Network structure of subsampled TCR repertoire of a male subject for DP, CD8, CD4 Teff and CD4 Treg SP (from left to right). Each point on the graph represents a CDR3aa. **(B-C)** Comparison of the proportion of linked sequences **(B)** and network density **(C)** between male and female samples, in DP (N = 22), CD8 (N = 23), CD4 Teff (N = 24) and CD4 Treg (N = 14). Each point on the graph represents the median value from 100 subsampling iterations for each sample. Statistical analysis using the Wilcoxon test revealed no significant sex differences for these two metrics (p > 0.05). Males are depicted in violet; females in orange.

### Similar network structure of thymic TCR repertoires in males and females

Next, we evaluated whether thymic selection in males and females is directed towards similar or distinct sequences. For each TCR repertoire, CDR3aa sequences are connected if they differ by one aa, i.e. have a Levenshtein distance of =1 (**Figure 1**). Such linked sequences have a high probability of sharing the same specificities (30). A random downsampling process to the smallest sample was performed to compare samples of comparable sizes. This process was repeated 100 times, and the results are detailed in the **Supplementary Table 1**. The shapes of the network are depicted in **Figure 6A**.

To analyze these clusterings with greater precision, we used various metrics. When comparing the number of clustered sequences in each individual network between males and females, no significant differences were observed for any cell type in either chain (**Figure 6B**). Similarly, the degree of sequence similarity, as indicated by the density of edges between sequences within an individual’s TCR repertoire, was comparable between males and females (**Figure 6C**). These findings indicate that there are no significant sex-specific differences in the similarity of CDR3aa sequences, suggesting that males and females produce CDR3aa sequences with the same degree of variability within their TCR repertoires.

### Absence of sex-specific CDR3aa motifs in thymic TCR repertoires of males and females

We then sought to determine whether we could detect CDR3aa sequence motifs that were more prevalent in one group compared to the other, in order to evaluate whether the selection of CDR3aa was biased towards particular sequences differently between males and females (**Figure 1**). To identify sequences with shared characteristics that might indicate a probable common specificity among individuals of the same group, the analysis focused exclusively on CDR3aa sequences of the TRB, as this region has been shown to have a greater influence on TCR specificity (31,32). Local motifs were defined using Gliph2 (33), which are a sequences of three to five aa, as well as global motifs, which are sequences of more than three aa where, at one position, an aa can be substituted by another aa if it has a positive BLOSUM62 score (34). Importantly, only motifs containing public CDR3aa sequences shared by at least two individuals were retained, thereby excluding private or individual-specific sequences from the analysis.

We identified several hundred motifs that were differentially expressed between males and females for each cell subpopulation. A greater number of motifs were specific to females, ranging from 426 motifs for DP to 278 motifs for CD4 Tregs, compared to males, ranging from 328 motifs for CD4 Teffs to 34 motifs for CD4 Tregs (**Figure 7A**). The identified motifs are mainly local motifs and largely confined to their respective sex group, indicating that motifs identified in females are predominantly found in females, and vice versa (**Supplemental Figure 10-13**).

**Figure 7:**
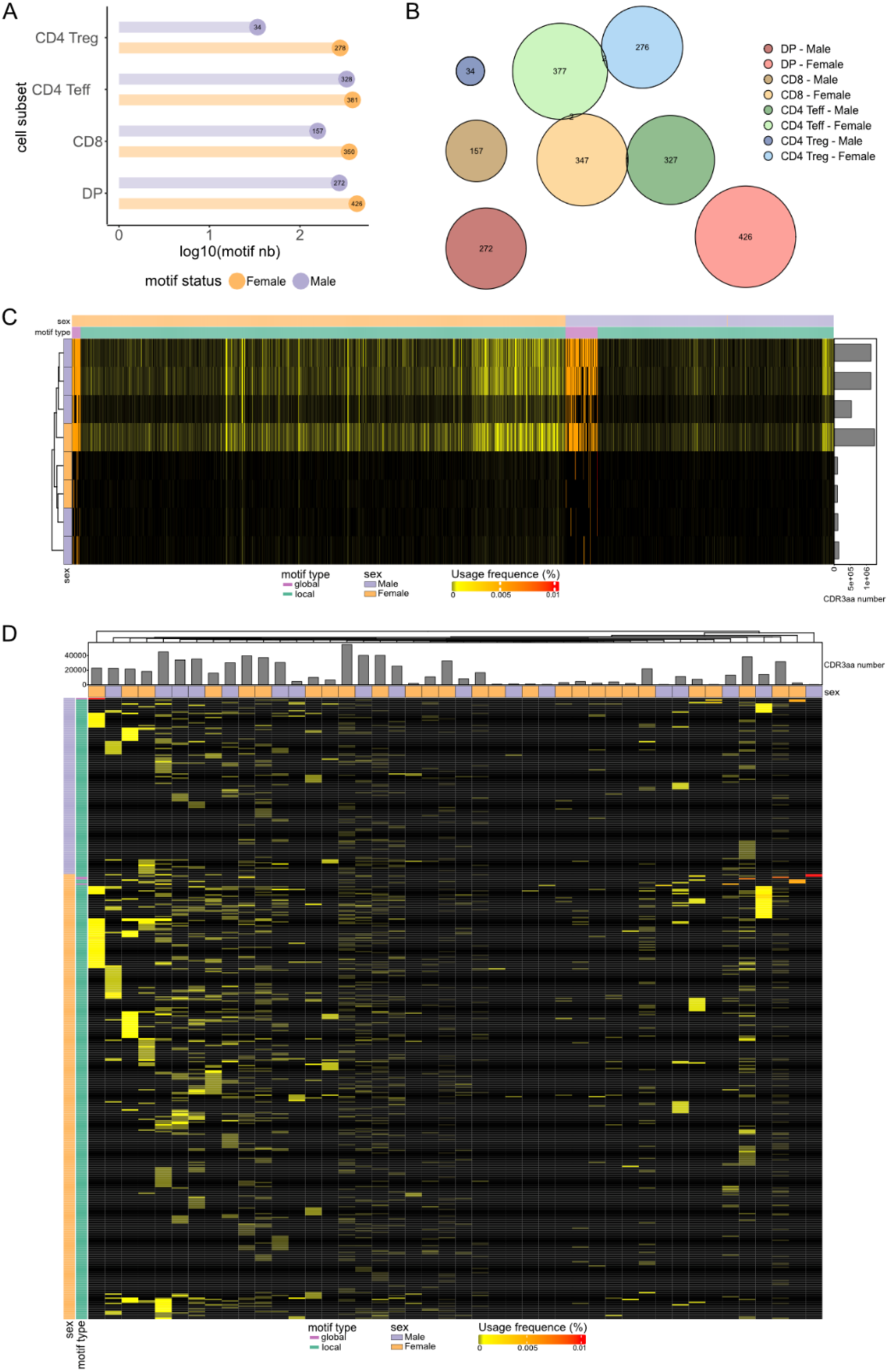
Thymic TRB TCR sex-associated motifs. Different structural motifs found differentially expressed between males and females in our dataset. We distinguish local motifs as distinct aa sequences, and global motifs as motif regions with one variable aa position maintaining a BLOSUM62 score of ≥ 0. **(A)** Number of male and female associated motifs by cell subset. **(B)** Euler diagram illustrating the distribution and overlap between all sex-associated motifs. The numbers indicate the number of motifs in overlap zones. **(C-D)** Validation of these sex-associated TRB CDR3aa motifs. The following heatmap illustrates the usage of all the TRB CDR3aa motif in the external thymic pediatric dataset (35,36) **(C)** and those of TRB CD8 in the peripheral dataset **(D)**. Sex and total CDR3aa number are depicted by sample. Males are depicted in violet and females in orange then local motifs in blue-green and global motifs in magenta.

As illustrated in **Figure 7B**, we observed minimal motif overlap between different cell subtypes. Only two female associated motifs overlapped: one between CD4 Teffs and CD4 Tregs, and another between CD4 Teffs and CD8 cells (**Figure 7B**). In contrast, no overlap was found between male-associated motifs (**Figure 7B**). Furthermore, it was observed that there was an overlap between one motif associated with female CD8 cells and a motif associated with male CD4 Teff cells (**Figure 7B**).

We then proceeded to evaluate the usage of the differentially expressed motifs in external datasets. There is no publicly available thymic TCR dataset that reports repertoires according to cell subsets. We thus tested these motifs on TCRs from pediatric bulk thymocytes, which contains TCR data from children aged between two and eight months, with a male-to-female ratio of 5:3 (35,36). We calculated the usage of these differentially expressed motifs in each sample of this dataset. We were unable to separate individuals by sex using these motif usages (**Figure 7C**).

We then evaluated these motifs using a peripheral blood TCR dataset from healthy individuals, where CD8, CD4 Teff, and CD4 Treg cells had been sorted. The usage of these motifs could not distinguish males from females across all cell subsets (**Figure 7D** and **Supplemental Figure 14**).

### Sex-biased enrichment of TCRs associated with autoimmune diseases and bacterial antigens

We sought to identify the usage of TCRs with known specificity as represented in the IEDB, Mc-PAS and VDJdb public databases (37–39). We compiled the TCRs from these three databases, retaining only those with high sequence reliability scores (see Materials and Methods). We focused on TRB sequences exclusively, due to their greater representation in the databases (31,40). We identified 55,368 unique TRB CDR3aa sequences with high reliability and specificity assignment scores for at least one specificity.

We classified the TCR specificities based on the antigen they recognize: viral or bacterial peptides, or human peptides overexpressed in cancers, autoimmune diseases (AIDs), or neither of these diseases (**Supplemental Figure 15A**). We sought to identify the TRB CDR3aa sequences in our thymocyte dataset. Depending on the individuals and cell subtypes, they represented between 0.82% and 3.58% of the TRB repertoires.

We observed that unique TRB CDR3aa sequences specific to self-antigens not associated with pathologies and those associated with cancers, are proportionally more represented in our thymic TCR dataset than in the pooled reference database, in both females and males (**Supplemental Figure 15B**). This pattern was consistent across all cell subtypes (**Supplemental Figure 15B**). These comparisons to the reference database are presented for descriptive purposes only, as differences between an experimental thymic repertoire and a curated database are expected given the structure of the reference resource. We then compared the distribution of these CDR3aa between males and females for each cell subtype. Strikingly, in the TCR repertoires with one specificity only, we observed a significantly higher proportion of unique TRB CDR3aa sequences associated with AIDs in female CD8 SP cells compared to male CD8 SP cells (**Figure 8A**). Furthermore, in the specific TCR repertoires only, there was a trend towards a higher proportion of thymic sequences with specificity associated to bacterial compounds in female CD8 SP cells compared to males, and an opposite trend was observed for CD4 Tregs SP (**Figure 8A**). These differences observed in CD8 SP cells persisted when examining the usage of these sequences (i.e. cumulative usage frequency), with significantly higher usage of sequences with specificity associated to bacterial compounds and those associated with AIDs in females compared to males (**Figure 8B**). Of particular note are the donor-resolved analyses (**Supplementary Figures 15B and 16**), in which each individual is represented separately. These confirm that these patterns are not driven by a single donor. Donor age had no significant effect on the sex-biased enrichment of self-and bacteria-specific TRB CDR3 sequences in CD8 SP and CD4 Treg SP cells (p ≥ 0.52), indicating that the observed differences are age-independent (**Supplemental Figure 17**). In addition, we investigated for the presence of polyspecific TCRs, which are capable of recognizing multiple antigens from different organisms (41–43). These sequences were found to be enriched across all cell subtypes, in comparison to their representation in the reference database (**Supplemental Figure 15C**). However, no significant differences were observed in the proportion or usage of these polyspecific sequences between males and females. Interestingly, an inverse usage trend was observed between CD8 and CD4 Treg SP cells: females exhibited higher usage of these sequences in the CD8 SP repertoire but a lower usage in the CD4 Treg SP repertoire compared to males (**Figure 8C-8D**).

**Figure 8:**
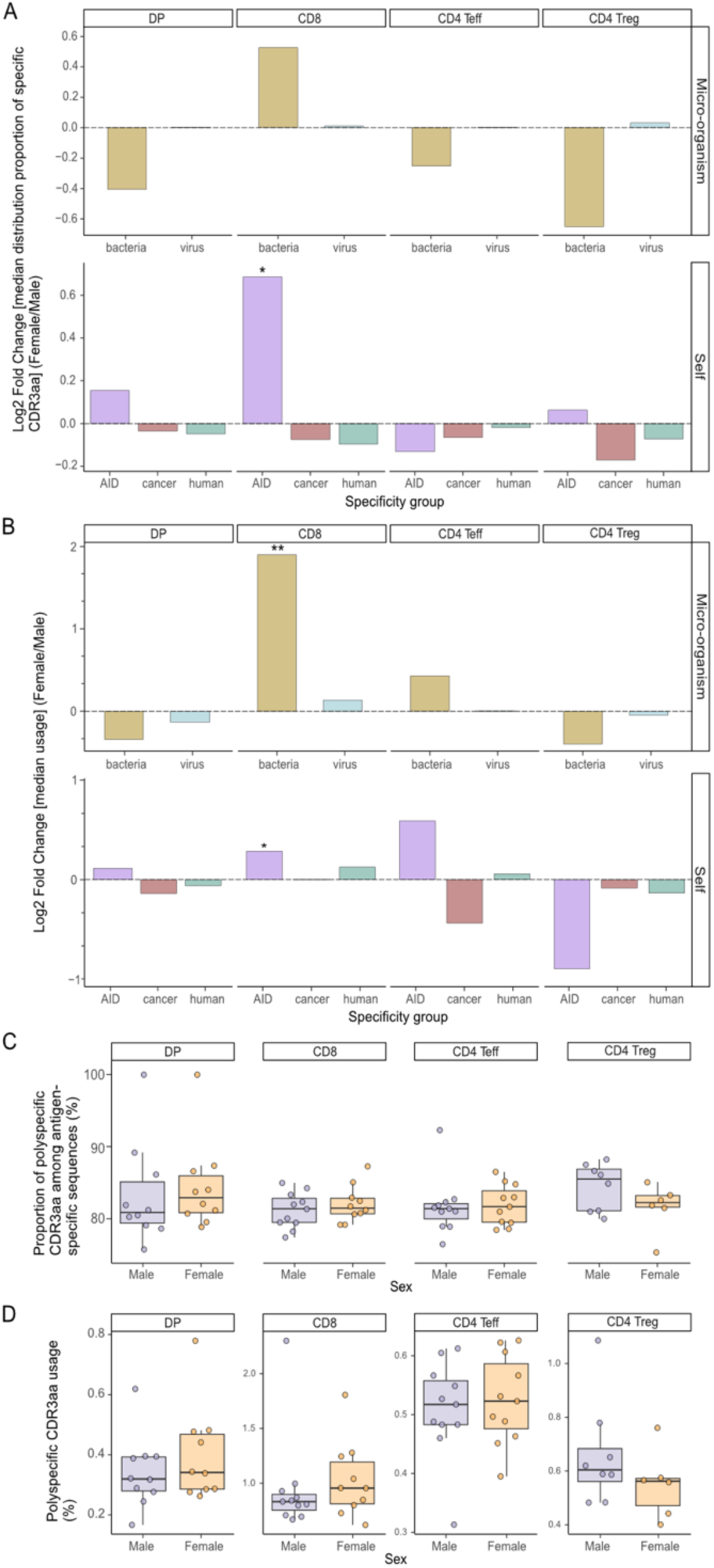
Sex-biased enrichment of TCRs specific for known antigens. From a pooled and curated database, an exact match with this database infers the specificity of TRB CDR3aa of our thymic dataset. Many specificity groups are defined according to the nature of the antigen peptide targeted (bacteria, virus, autoimmune disease [AID], cancer and self-peptide no associated to disease [human]). This analysis compares the distributions of the proportion of unique TRB CDR3aa sequences with a specific specificity **(A)** and their usage **(B)** between females and males across cell subtypes, using the log2 fold change of the median values (females over males), following each specificity group. These groups of specificity are additionally classified as microorganism in the top section (bacteria in gold and virus in light blue) and self at the bottom (AID in magenta, cancer in red, human in blue-green). Polyspecific CDR3aa are defined here as CDR3aa capable of recognizing multiple antigens from different organisms (for no self-antigens) or from different specificity groups (e.g. categorized microorganisms, categorized self-antigens, allergens…). The proportion of polyspecific CDR3aa among antigen-specific sequences **(C)** and their usage **(D)** is compared between males and females, in DP (N = 22), CD8 (N = 23), CD4 Teff (N = 24) and CD4 Treg (N = 14) cells. Stars indicate statistical differences between males and females based on the p-value of the Wilcoxon test (*: p < 0.05, **: p < 0.01). Males are depicted in violet and females in orange.

In conclusion, our analysis of TCR specificity has identified sex-specific differences in the thymic selection of CD8 effector T cells associated to autoimmune and bacterial antigens that are overrepresented in female versus males.

## Discussion

It is noteworthy that there is a paucity of research on potential differences in the TCR repertoire between the sexes, despite the much higher prevalence of autoimmune diseases in females, a topic that remains to be fully elucidated. To explore the hypothesis that the generation and/or the selection of the TCR repertoires may be different between sexes requires, it is necessary to study the TCR repertoire where they are formed and selected, in the thymus. We thus generated a unique and valuable dataset comprising thymic samples collected from deceased organ donors of various ages, and young children who have undergone cardiac surgery. Firstly, we separated the various thymocytes populations in order to study both TCR repertoire generation at the level of DP cells, and repertoire selection at the level of CD8, CD4 Teff and Treg SP cells. We performed bulk sequencing of the TCRs from these cells, as this is the only method that yet generates enough sequences for sensitive comparison at this time. Single-cell sequencing is an analyzing technique that it is limited in its scope. It is only capable of analyzing a few thousand cells and would not have enabled the analyses that can be performed with hundreds of thousands of cells.

### TCR generation

Our analyses initially revealed similarities rather than dissimilarities in TCR generation. Our detailed analyses revealed no significant sex differences in relation to the majority of the examined variables when comparing males and females DP cells’ repertoires, except for differences in the usage of small number of V and J genes from both TCR chains. However, these minor discrepancies are likely attributable to random variation, considering the extensive number of comparisons conducted. This finding is further supported by the observation that there are no significant differences in the usage of TRBV-TRBJ and TRAV-TRAJ gene associations between males and females. We also observed no differences in diversity indexes in DP cells’ repertories between males and females. Altogether, the overall combinatorial diversity of the TCR repertoire remains comparable between sexes.

A detailed analysis of the CDR3aa region in DP cells showed that some hydrophobic aa in the FG loop region are differently found for both TCR chains. However, no significant sex-based differences were found in the usage of hydrophobic aa at the critical p109 and p110 positions in TRB that have been described as influencing self-antigen recognition due to the impact of hydrophobic aa on TCR self-reactivity (26–28). These findings suggest that the subtle sex-based differences in the FG loop’s aa usage are minor and do not significantly impact the overall characteristics of the CDR3 region in a sex-dependent manner.

The Pgen analysis of DP cells revealed only minimal differences in the distribution of Pgen between males and females, which were similar to those observed in control groups. This indicates that the mechanisms governing TCR generation are largely consistent across sexes, with only minor, likely random, deviations.

Additionally, although we identified TRB CDR3aa sequence motifs that were differentially expressed between males and females at the DP cell stage in our dataset, these motifs could not reliably differentiate individuals by sex in external datasets. This indicates that they are more dataset-specific than sex-specific. This issue may also be explained by the different characteristics of our data and that of the other dataset. For example, in the Arstila dataset, the analyses were performed on unsorted cells from young subjects. Younger individuals tend to have a more diverse and richer TCR repertoire, with greater repertoire sharing in the periphery, which could potentially obscure sex-specific patterns (16,44). This suggests that these motifs might be context-specific or influenced by other factors such as age, genetic background, or environmental exposures. Moreover, we did not observe any sex-specific differences in CDR3 TRB specificity usage in DP cells’ repertoires. In summary, our results from DP cells analyses could not identify any relevant sex differences in TCR generation.

### TCR selection

We identified three genes that show differential usage between males and females. Whilst it is possible that some of these differences are merely random, resulting from multiple comparisons, the preferential use of TRBV6-5 in females has previously been observed in the peripheral TCR repertoire (45). Notably, this gene is overrepresented in female lupus patients compared to their controls (51,52). Although the preferential usage of TRBV6-5 in females is consistent with previous peripheral observations, external validation of this thymic bias remains challenging. Publicly available TCR datasets rarely provide (i) sorted thymocyte subsets, (ii) balanced sex representation, and (iii) sufficient sequencing depth within each subset to allow reliable comparison of gene usage patterns. Moreover, differences in tissue origin (peripheral blood versus thymus), age distribution, and technical protocols may substantially influence TRBV usage frequencies. For these reasons, direct replication in independent thymic cohorts is currently not feasible. Our permutation-based internal control analysis was therefore implemented to ensure that the observed signal was not attributable to random donor grouping or individual-level variation.

Our most striking observation relates to significant sex-specific differences in inferred specificities of CD8 T cells, with a biased CD8 selection process in the thymus favoring sequences associated with autoimmune diseases and bacterial antigens in females. The fact that this bias does not affect DP cells indicates that it is related to TCR selection rather than generation. Its relevance is strongly supported by the fact that (i) there is no such bias for other self-antigens that are not associated with autoimmunity and (ii) the bias affects both the effectors and regulators of autoimmunity with a directionality compatible with the increased prevalence of autoimmune disease in females. TCRs associated with autoimmunity-related self-antigens have been observed to be increased in CD8 effector T cells, while decreased levels have been seen in CD4 Tregs. It should be noted that this bias also affects certain bacterial antigens, though not viral antigens. This phenomenon could be explained by the fact that bacterial antigens express numerous mimotopes of self-antigens, and are known to contribute to the shaping of the TCR repertoire. The observed bias towards bacterial antigens in female CD8 T cells could also provide more effector cells specific for bacterial antigens that mimic self-antigens, contributing to autoimmune disease development by molecular mimicry (48). In this line, antigens linked to celiac disease (CeD) and type 1 diabetes (T1D) are highly represented in the database and these two AIDs have been linked to bacterial components that may mimic self-antigens (49–53). As is well-established, the gut microbiota influences thymic development of T cells (54). Sexual hormones could influence this biased selection by affecting the composition of the gut microbiota (55) and/or by affecting the selection of the TCRs specific for these antigens. In addition to these specific patterns, we observed a significant increase in the usage of hydrophobic amino acids at the central p109 position of the TRB CDR3aa in female CD8 thymocytes, with similar trend at p110. Hydrophobic residues at these positions have been implicated in enhanced self-reactivity and cross-reactivity of CD8 T cells (26–28), suggesting that female CD8 thymocytes may be slightly enriched for TCRs with intrinsically higher potential for recognizing self-like or cross-reactive peptide MHC complexes. It is noteworthy that the positional hydrophobicity of CD8 T cells CDR3s is consistent with the over-representation of sequences annotated as recognizing autoimmune-associated self-antigens, pointing to a more autoreactive TCR repertoire.

Finally, we have recently described polyspecific TCRs that can respond to multiple unrelated viral antigens and hypothesized their possible involvement in autoimmunity (43). We defined a CDR3 as being polyspecific if the TCRs containing it have been experimentally associated with epitopes from at least two different antigen origins. We observed a higher usage of polyspecific sequences in the CD8 SP repertoire of females, although this was not statistically significant. In contrast, the frequency of polyspecific TCR sequence usage was lower in female CD4 Tregs SP cells. These observations, which could be indicative of improved anti-infectious responses and worse autoimmune responses in women, warrant further exploration.

These inferences rely on a harmonized specificity compendium combining McPAS-TCR, IEDB and VDJdb, which provide complementary and only partially overlapping landscapes (pathology-enriched versus predominantly viral specificities). Although there is limited numerical overlap between our thymic repertoires and annotated sequences is limited, this is to be expected given that public databases currently capture only a small fraction of the potential TCR specificity space. In this context, the sex-biased enrichment of sequences annotated as autoimmune-and bacteria-associated should be viewed as a robust signal emerging from a very sparse annotation space. However, it should still be interpreted as hypothesis-generating rather than exhaustive.

To further evaluate the robustness of our findings, we performed additional analyses to assess whether donor age could confound the observed sex-specific differences. A potential limitation of our study is the relatively wide age range of donors. However, the absence of distinct age-related clustering in repertoire features, the lack of age impact on CD8 and CD4 Treg bacterial-specific TRB usage and CD8 and CD4 Treg self-associated to autoimmune disease-specific TRB usage patterns (Supplemental Figure 17), argue against age as a confounding factor. These results support the hypothesis of an intrinsic sex bias in thymic TCR selection rather than an age-related effect.

While our findings provide valuable insights into sex-specific differences in thymic selection of TCR repertoires, our TCR specificity analyses were limited. We can confirm that our study focused exclusively on the TRB CDR3aa region of the TCR, which is widely acknowledged as the most critical element in defining TCR specificity (31,32) while both the TRA and the TRB chains contribute to TCR specificity (31,32). This limitation is intrinsic to the limited number of TRA with assigned specificity in databases and to the fact that, given the nature of our studies, there is a need to study large repertoires, which is currently not feasible by single cell TCR sequencing. However, as methods for inferring specificity continue to evolve, they will likely allow more refined analyses of our dataset in the future (56). Furthermore, the continuous development of the TCR database should enhance these analyses.

Despite the relatively small size of our cohort, which was outbred, we believe that this design is essential for capturing sex-associated differences. Inbred models would reduce inter-individual variability, but they would not reflect the complexity of human thymic selection, nor the interplay between sex and genetic diversity. Despite the modest sample size, the consistency of the observed patterns across the analytical approaches supports the robustness of our conclusions.

Altogether, our analyses of TCR specificity identified a biased selection in females towards sequences associated with autoimmune diseases, supporting the idea that early central tolerance mechanisms may represent one layer contributing to sex differences in autoimmune susceptibility, in interaction with peripheral and environmental factors. Future research should focus on validating these observations in relevant independent datasets, which are currently lacking.

## Methods

### Samples

Thymus samples were obtained from twenty-two organ donors aged between 73 days and 64 years, without any specific pathologies (**Figure 1**). Individual partial HLA typing is detailed in **Supplementary Table 2**. Adult samples were collected post-surgery at the Cardiac Surgery Departments of many hospitals in France, following approval by the Agence de Biomédecine and the Ministry of Research (#PFS14-009). Pediatric samples were obtained following total thymectomy during cardiac surgeries at Necker Hospital. The ratio of male to female across donors is maintained at 1:1, although it varies between 1.1:1 and 1.33:1 depending on the cell subtypes analyzed. The age distributions of male and female donors were compared using two-sample Kolmogorov–Smirnov tests, with no significant differences observed across thymic subsets (p > 0.46), indicating no age-related confounding (**Supplementary Figure 1**).

### Thymocyte isolation and RNA extraction

Thymic cell suspensions were prepared through automated tissue dissociation using gentleMACS dissociators (Miltenyi^®^) or by mechanical disruption, followed by filtration through a 70-µm nylon mesh. The cells were stained with anti-CD3 (AF700), anti-CD4 (APC), anti-CD8 (FITC), and for some samples, anti-CD25 (PE) antibodies. Cell sorting was performed by fluorescent activated cell sorting (FACS) on a Becton Dickinson FACSAriaII, achieving a purity of >85%. The sorted populations included DP CD3+ (CD3+CD4+CD8+), SP CD8+ (CD3+CD4-CD8+), and SP CD4+ (CD3+CD4+CD8-). A further separation was made into SP CD4 Teff (CD3+CD4+CD8-CD25-) and SP CD4 Treg (CD3+CD4+CD8-CD25+) for sixteen of the samples (**Figure 1**). For the sorted SP CD4, the Treg percentage varied between 5.8% and 16% across samples, with an average of 9.63%. Six samples of SP CD4 samples were not sorted into Teffs and Tregs, we thus categorized them as SP CD4 Teffs, assuming a purity of Teffs within the total SP CD4 of around 90%. Please refer to the detailed sample metadata in **Supplementary Table 3** of the supplementary material. This includes donor IDs and the number of sorted cells per thymic subset for each donor.

The RNA was isolated using the RNAqueous extraction kit (Invitrogen^®^), in accordance with the manufacturer’s protocol. The RNA samples were quantified and the integrity of each sample was determined using a Nanodrop (Thermo Fisher) or a Tapestation 4200 (Agilent, RNA ScreenTape).

### TCR library preparation and sequencing

TCR library preparation and sequencing were performed as described by Barennes et al. (57). Briefly, RNA was processed using the SMARTer Human TCR a/b profiling v1 kit (Takarabio), in accordance with the manufacturer’s protocol. Amplicons were then purified using AMPure XP beads (Beckman Coulter) and their quantity was determined. The integrity of the amplicons was then assessed using the 2100 Bioanalyzer System (Agilent, DNA 1000 kit) or Tapestation 4200 (Agilent, D1000 screentape). Bulk Next-Generation Sequencing was performed using either a Hiseq 2500 (Illumina) with the SR-300 protocol plus 10% PhiX on the LIGAN-PM Genomics platform (Lille, France), or a Novaseq 6000 (Illumina) with the PE-250 protocol plus 10% PhiX on the LIGAN-PM Genomics platform or ICM platform (Paris, France).

### Data processing

The raw FASTQ/FASTA files were aligned to TRA and TRB using MiXCR (v3.0.13, RNA-seq parameters), a software solution that is able to correct for both sequencing and PCR errors (58). Samples with fewer than 1,000 CDR3aa sequences, an imbalanced TRA/TRB ratio (<1/9), or a low sequence count relative to sorted cell counts (< 0.9 ratio) were removed. Sequences with CDR3aa lengths outside 6-23 amino acids were also discarded. To standardize clonotype counts and the number of unique clonotypes across sequencing techniques, the initial counts for each clonotype were reduced by 1 to exclude those with counts of 0 (singletons).

Furthermore, due to the high similarity between certain TRBV sequences, which has the potential to influence the analysis depending on the sequencing method used, TRBV06-2 and TRBV06-3 were merged as TRBV06-2/3, and TRBV12-3 and TRBV12-4 were combined as TRBV12-3/4 (59). Please refer to the summary table provided in the supplementary material (**Supplementary Table 3**) for a comprehensive overview of the total number of TCR sequences retained after preprocessing, as well as the corresponding number of unique clonotypes, for each donor, chain and thymic subset.

### Gene usage analysis

For both TRA and TRB chains across all cell subtypes, we analyzed the V and J gene usage and those of the VJ gene associations (**Figure 1**).

The usage of gene is calculated as follows: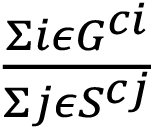

Where:

- Σ𝑖𝜖𝐺*^Ci^* represents the total number of counts for clonotypes associated with the specified gene or gene association.
- Σ𝑗𝜖𝑆*^cj^* represents the total sum of the counts of all clonotypes in the sample.

In this notation, 𝐺 is the set of clonotypes using the interested gene, 𝑆 is the set of all clonotypes in the sample, and c*_i_* and c*_j_* are the counts of the clonotypes *i* and *j*, respectively. The usage of the V gene and the VJ gene combinations was investigated through dimensional reduction using Principal Component Analysis (PCA) (MASS package v7.3-60, factoMineR package v2.8, factoextra package v1.0.7). Confidence ellipses were represented with a 5% alpha risk. Furthermore, for VJ gene combinations usage, a heatmap was constructed using the Jensen-Shannon Divergence (JSD) distance to measure the distance between samples based on their gene usage distribution (complexHeatmap package v2.16.0, philentropy package v0.7.0). Hierarchical clustering was performed using the Euclidean distance and the complete linkage method.

### Diversity analysis

For the TCR repertoire diversity analysis, to correct for sequencing depth bias and differences in cellular richness between individuals, rarefactions were performed fifty times for each sample, using X clonotypes, where X represents the effective diversity number (60).

The effective diversity was calculated as the exponential of the Shannon index (𝑒^!^) (61). Rényi indices (11 indices from 0 to Infinity) were calculated for each rarefaction (62).

The median Shannon index, Simpson index and Berger-Parker indices were then compared between the sexes.

### Distribution of CDR3aa length analysis

The calculation of the usage of all CDR3 by the amino acid length as follows: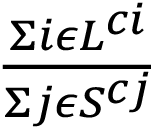

Where

- Σ𝑖𝜖𝐿*^ci^* represents the sum of the counts of the CDR3 characterized by the amino acid length of interest.
- Σ𝑗𝜖𝑆*^cj^* represents the total sum of counts of all CDR3s per sample.

In this notation, 𝐿 is the set of CDR3s characterized by the amino acid length of interest, 𝑆 is the set of all CDR3s in the sample, and c*_i_* and c*_j_* are the counts of the CDR3s *i* and *j*, respectively.

### Composition usage of amino acid within the CDR3 region

All CDR3aa sequences were aligned using the IMGT convention (positions p104-p118) (63).

Three complementary scales were used in the study. Firstly, sequence level amino acid composition. Within the CDR3 loop region (IMGT positions p108-p114), involved in MHC-peptide complex interaction (63), amino acid usage at the sequence level was quantified for CDR3aa sequences 9-21 aa long in both TRA and TRB chains. For each sequence, the percentage usage of a given amino acid was calculated as follows:

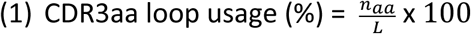

Where 𝑛*_aa_* is the number of occurrences of the amino acid of interest in the CDR3 loop, and *L* denotes the total length of the CDR3aa sequence. Furthermore, amino acids were categorized into three hydropathy classifications (neutral, hydrophobic, and hydrophilic) using the IMGT classification that takes into account the Kyte-Doolittle hydropathy index (25,64)

Secondly, the position-specific amino acid usage at p109 and p110. To assess positional composition, for each individual, CDR3aa usage is examined at IMGT positions p109 and p110, corresponding to p6-p7 positions described by Stadinsky et al. and Khosravi-Maharlooei et al., which hydrophobic amino acids at these two central positions have been implicated in modulating TCR self-reactivity and cross-reactivity (26–28). For a hydrophobic amino acid ℎ (excluding alanine, due to its weak hydrophobicity) at position 𝑝, usage was defined:

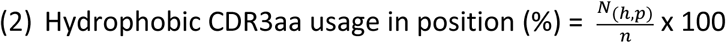

Where 𝑛 is the total number of unique CDR3 sequences and 𝑁_(*h,p*)_ the number of unique CDR3 sequences carrying hydrophobic amino acid 𝑎 at position 𝑝. This analysis quantifies amino acid composition at fixed positions independently of clonotype frequency.

### Probability of generation analysis

The probability of generation analysis of DP and CD8 cell subtypes consisted initially in the creation of a generative model of V(D)J recombination using 400,000 randomly selected non-productive sequences from all samples for both TRA and TRB chains, with the IGOR tool (65). The generation probabilities (Pgen) for all sequences in the dataset were subsequently calculated using the OLGA tool, based on the respective generative models for each cell subtype and TCR chain (66). Comparisons of Pgen values at the nucleotide level between males and females were performed. To ensure statistical robustness, control groups with equivalent male and female sample numbers were created by permuting the samples into two groups 20,000 times.

### TCR network structure by sequence similarity

For each sample, 100 random sub-samplings of CDR3aa sequences were performed, using the minimum CDR3aa count per cell subtype (refer to **Supplementary Table 1**). Two CDR3aa were connected if their Levenshtein distance was = 1 (**Figure 1**) (67). Network analysis focused on two metrics: the proportion of connected sequences and network density, defined as the ratio of actual connections to all possible connections. Median values from 100 subsampling iterations were then analyzed. Levenshtein distances were computed with the stringdist package (v0.9.10), networks were visualized using Cytoscape (v3.8.2), and network density was calculated with the igraph package (v1.5.1).

### TRB CDR3aa motif search

Two types of structural motifs were searched: local motifs (strict sequential motifs) and global motifs (sequences with substitutions preserving a positive BLOSUM62 score) (Figure 1).

The selection of enriched motifs was carried out using the gliph2 function with the turboGliph package (v0.99.2), with specific parameters defined as follows:

-The input sequences were from the sex group of interest (male or female), with reference sequences matched for TRBV gene usage.

-It is imperative that CDR3aa sequences are longer than eight aa.

-Local motif lengths were restricted to 3-5 aa.

-Only motifs consisting of more than two unique CDR3aa were conserved.

-Local motif significance of Fisher’s test has not been boosted.

Additional filters were applied so that: (i) the inclusion of public CDR3aa sequences is a prerequisite for a motif, with these sequences shared by at least two individuals, (ii) a significant enrichment is required, as determined by Fisher’s test with a p-value p < 0.01 and (iii) a usage difference between groups of at least twofold is necessary, as evidenced by a Wilcoxon test with a p-value p < 0.05.

Motif validation was performed on an openly published pediatric thymic dataset from the Arstila team (35,36) consisting in a bulk TCR dataset from infants aged between seven days and eight months. Here, we removed one male subject, “Thymus A”, who was a twin of the “Thymus B” and “Thymus 1” neonate sample, in order to analyze height samples with a male-to-female ratio of 5:3. Further validation was conducted on sorted peripheral blood TCR repertoires (CD8, CD4 Teff and CD4 Treg) from seventy-five healthy volunteers aged from eighteen to eighty-four years (male-to-female ratio of 1.03:1), whose libraries were generated under the same conditions as those of our thymic dataset. They were recruited within the Transimmunom observational trial (NCT02466217) and HEALTHIL-2 (NCT03837093) (68). PBMCs from healthy individuals were isolated using Ficoll density gradient centrifugation and enriched for CD4+ T cells via EasySep™ Human CD4+ T Cell magnetic beads (Stemcell). The samples were then separated into two groups: effector T cells (CD4+ CD25-) and Tregs (CD4+ CD25+). This was done using EasySep™ Human Pan-CD25 magnetic beads (Stemcell). Purity was assessed by flow cytometry based on the expression of CD4 and FoxP3, with a purity threshold of >80%. From the frozen PBMC aliquots, CD8+ T cells (CD3+ CD8+) were sorted using a FACS ARIA II cell sorter.

#### TRB CDR3aa specificity analysis

We have established a curated and harmonized TCR specificity database by integrating data from three public TCR repositories: Mc-PAS, IEDB, and VDJdb (38,39,69) (data collected as of October 2023) (**Figure 1**). The harmonization procedure was carried out in accordance with the framework outlined in Jouannet et al. (70) and included:

-Uniform formatting of TRA/TRB TCR sequences and associated metadata,

-Consolidation of redundant entries (identical TRA/TRB TCR, epitope, organism, PubMed ID, and cell subset),

-Maintening separate entries when source databases are incompatible,

Reliability levels for sequences and specificity assignments were quantified using the Verified_score (VS) and Antigen-identification_score (AIS) as defined in (70). The VS, ranging from 0 to 2, reflects the concordance between calculated and curated CDR3 boundaries following the IEDB verification strategy (VS = 2: both TRA and TRB present and verified; VS = 1.1: only TRA verified; VS = 1.2: only TRB verified; VS = 0: no verified chain) and the AIS (0 to 5), which ranks the strength of antigen-identification methods (70).

In this study, we focused our analyses on high-reliability TRB sequences, defined as those with a verified TRB CDR3aa (VS ≥ 1.2) and an AIS corresponding to in vitro T cell stimulation with a pathogen, protein or peptide, or pMHC X-mer sorting (AIS > 3.2, excluding categories 4.1 and 4.2). This ensures that the specificity assignments relied on robust experimental evidence.

Specificity groups were categorized according to antigen class, encompassing viral, bacterial, yeast, parasitic and self-antigens along with antigens derived from wheat, plants or animals. Self-antigens were then annotated into three categories: autoimmune-associated antigens, cancer-associated antigens and self-antigens unrelated to disease. For autoimmune-associated antigens, assignments were based on explicit annotations present in McPAS-TCR,

IEDB and VDJdb. These databases identify celiac disease (CeD) and type 1 diabetes (T1D) related antigens, which were retained as such in the harmonized dataset. For cancer-associated antigens, the following classifications were used: (i) information present in the original databases and (ii) cross-referencing with external cancer antigen resources, including the Cancer Antigenic Peptide Database (de Duve Institute), the Human Protein Atlas and the cancer database caAtlas (71). Please refer to **Supplementary Table 4**, which provides a comprehensive list of all self-antigens present in the harmonized database. This table also includes their associated disease category and their mapping into the specificity groups used in the manuscript.

This harmonized database was used to identify exact matches with the CDR3aa sequences of our TRB sequences. The enrichment of CDR3aa sequences associated with a given specificity group in male and female thymic repertoires for each cell subtype compared to the specificity distribution in the database was tested. Furthermore, we performed specificity group distributions between male and female samples, as well as was the overall usage of these sequences across entire TCR repertoires.

Additionally, our research concentrated on polyspecific sequences defined as CDR3aa TRB sequences recognizing at least two distinct microbial species or belonging to multiple specificity groups for non-microbial antigens. In this study, polyspecificity is therefore considered at the level of broader antigenic categories rather than individual epitopes and thus reflects functional cross-reactivity patterns aggregated across studies, instead of implying molecular-level multi-epitope recognition. These polyspecific TRB CDR3aa sequences underwent the same statistical analyses as monofunctional sequences.

## Statistical analysis

For V-J gene usage, CDR3aa length usage, CDR3aa usage, network similarity and specificity analyses, the comparison between males and females was statistically analysed using the Wilcoxon test. The statistical tests were performed using the ggpubr library (v0.6.0) in R. The number of asterisks denotes the significance of the results: one asterisk indicates a p-value p ≤ 0.05, two asterisk indicate a p-value p ≤ 0.01, and so on until four asterisks, denoting a p-value p ≤ 0.0001. The comparisons of the Rényi curves and Pgen were compared using the Kolmogorov-Smirnov test. The Wilcoxon test was used to test the comparisons of the proportions of specific sequences in male and female subjects relative to those in the specificity database.

## Data availability

All raw sequencing data generated in this study are publicly available on NCBI under the BioProject accession PRJNA1379632 (https://www.ncbi.nlm.nih.gov/sra/PRJNA1379632).

## Author contributions

VQ and HV conducted all cell sorting. RNA extraction and TCR library preparation were performed by HV, VQ, LA, PB, NC, VD, JD, GF, KL and PS. HV, VQ, VM, LA, KL, ONT, MP, PS, AS, EMF and DK contributed valuable suggestions for experimental design and participated in discussions of the results. HV and CJ pooled and curated the reference specificity database. LA, CJ and VM contributed equally in this work. HV authored the manuscript, with VQ, VM, CA, JD, EMF and DK providing comments and critical reviews. DK conceptualized, supervised and provided funding for the study.

## Acknowledgment

We would like to thank Bruno Gouritin for his assistance with cell sorting and to Marie Surroque for her support with RNA quantification. During manuscript preparation, the authors used ChatGPT (OpenAI) solely to assist with language editing (syntax, grammar and phrasing). All ideas, analyses, interpretations and scientific reasoning are the authors’ own.

---

**Supplemental Figure 1:**
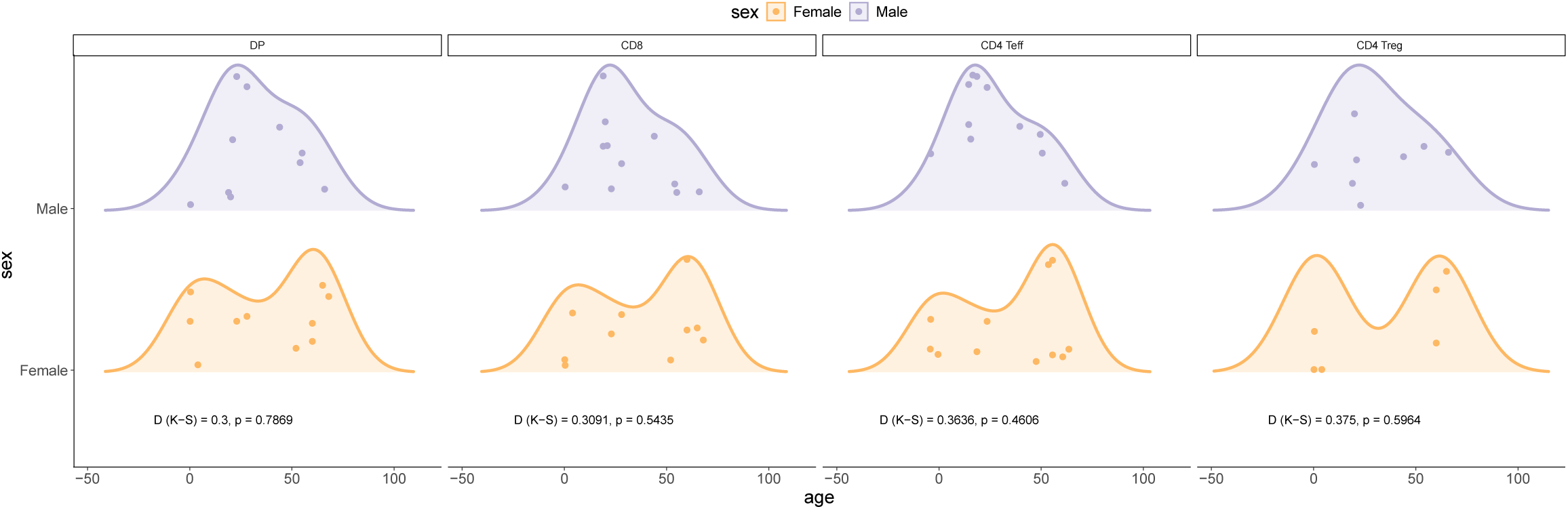
Age distribution of male and female donors is comparable across thymic subsets. Age of individual donors plotted separately for females (orange) and males (violet) for each thymic cell subset: double-positive (DP) thymocytes, CD8 single-positive (CD8), CD4 effector single-positive (CD4 Teff) and CD4 regulatory T-cell single-positive (CD4 Treg). For each subset, a two-sample Kolmogorov–Smirnov (K-S) test was performed; the D statistic and corresponding p-value are displayed below each panel. In all cases, no significant difference was detected, indicating balanced age distributions between sexes for every subset.

**Supplemental Figure 2:**
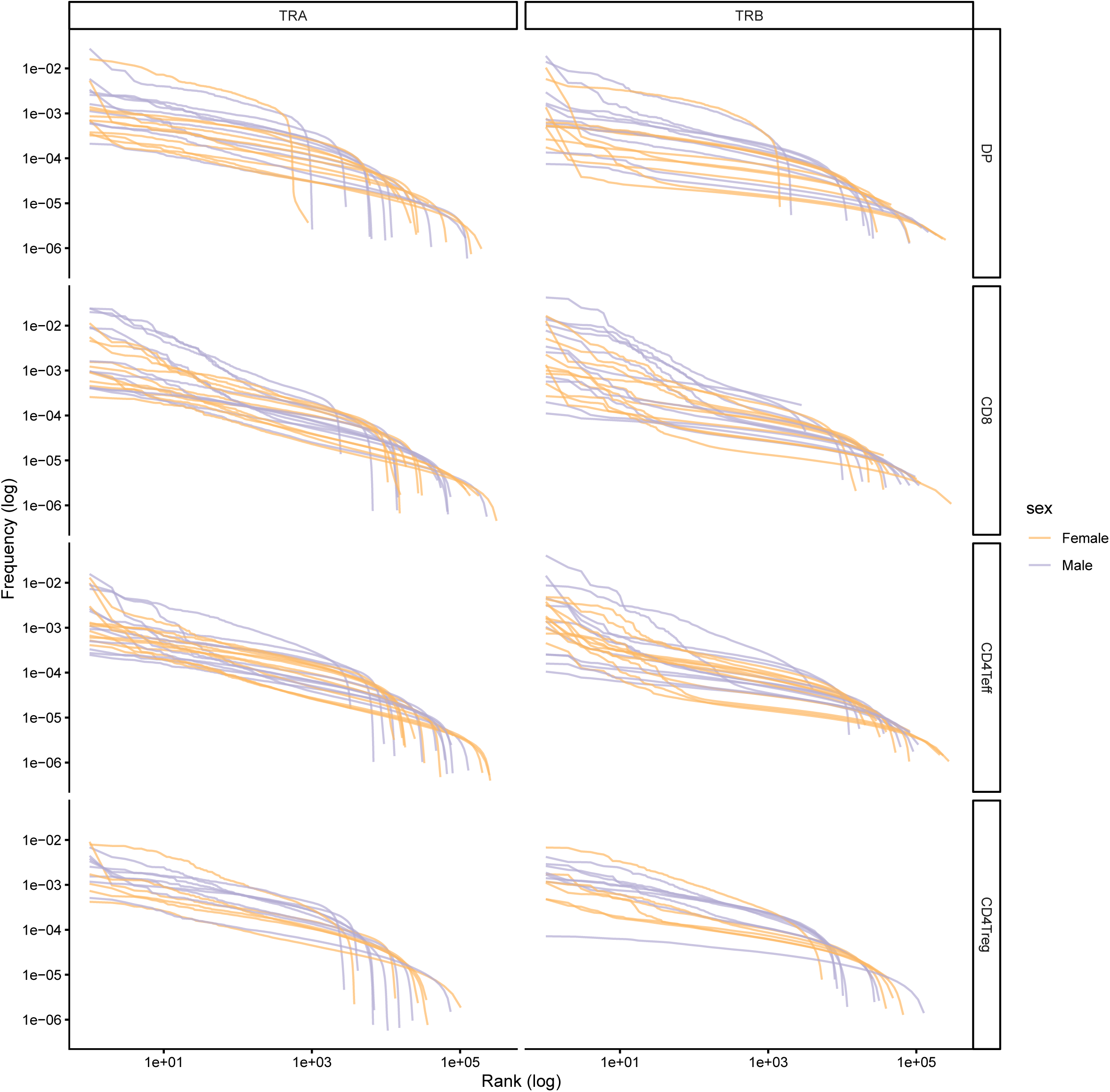
Rank–frequency distributions of thymic TRA and TRB clonotypes. Log–log rank–frequency plots of unique TRA (left) and TRB (right) clonotypes are shown for each donor and thymic subset (DP, CD8, CD4 Teff, CD4 Treg, from top to bottom). For each sample, clonotypes are ranked by decreasing abundance (x-axis, log scale), and their corresponding relative frequencies are plotted (y-axis, log scale). Curves display broadly comparable shapes across donors, with no obvious systematic differences between males and females, indicating similar clonotype abundance distributions and sampling depth across sexes.

**Supplemental Figure 3:**
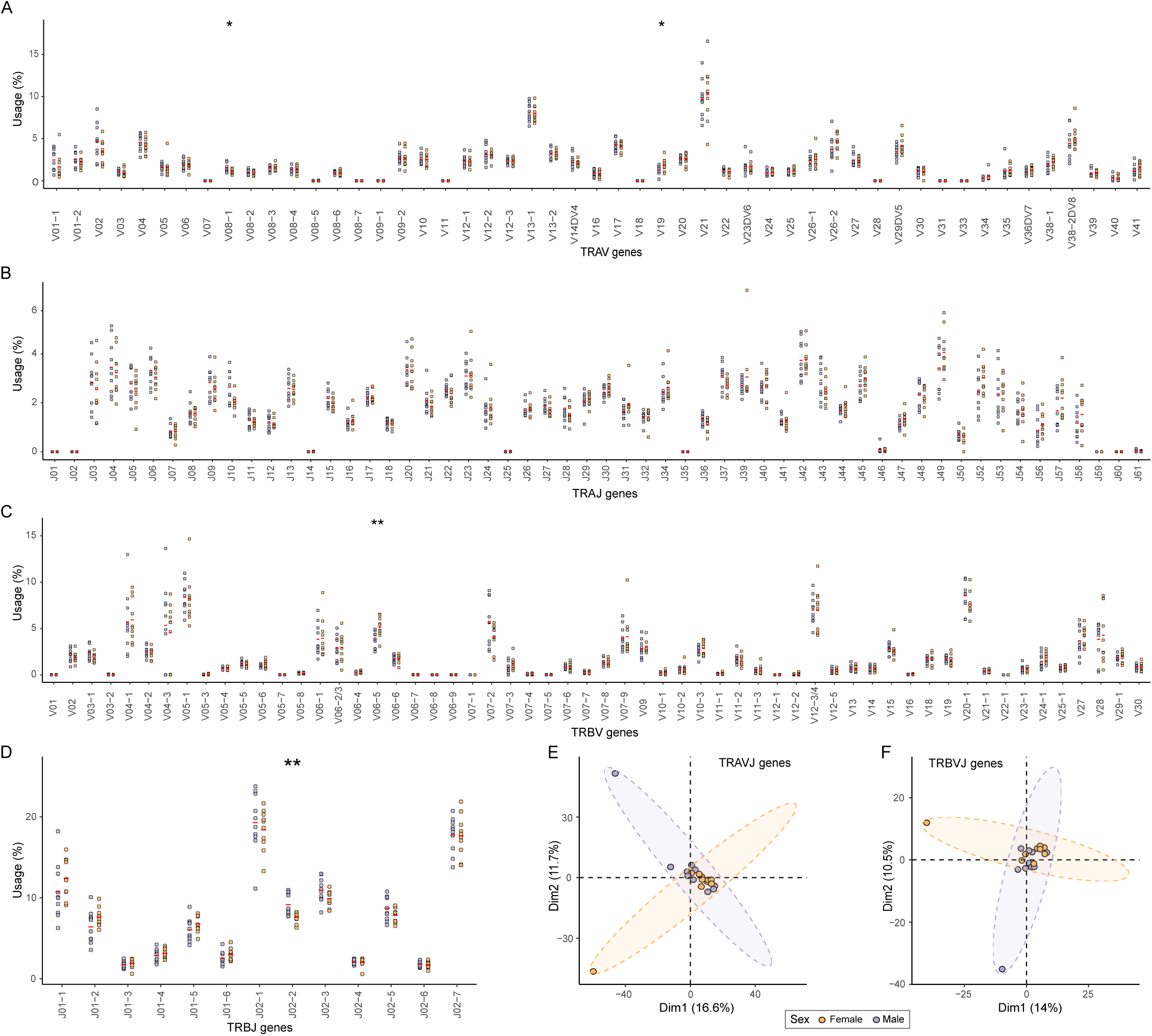
**Some differential V and J gene usage in TCR repertoires of DP cells between males and females**. **(A-D)** Frequencies of TRAV **(A)**, TRAJ **(B)**, TRBV **(C)**, and TRBJ **(D)** gene usage between males and females. Red lines represent the mean for each sex group. Statistical comparisons between groups were performed using the Wilcoxon test. Stars indicate statistical differences between males and females based on the p-value of the test (*: p < 0.05, **: p < 0.01). **(E-F)** Principal Component Analysis (PCA) derived from the distribution of the frequency of usage of TRAV-TRAJ **(E)** and TRBV-TRBJ **(F)** gene associations across sex groups between males and females. Each point represents an individual. Ellipses show 95% confidence intervals. Males are depicted in violet and females in orange.

**Supplemental Figure 4:**
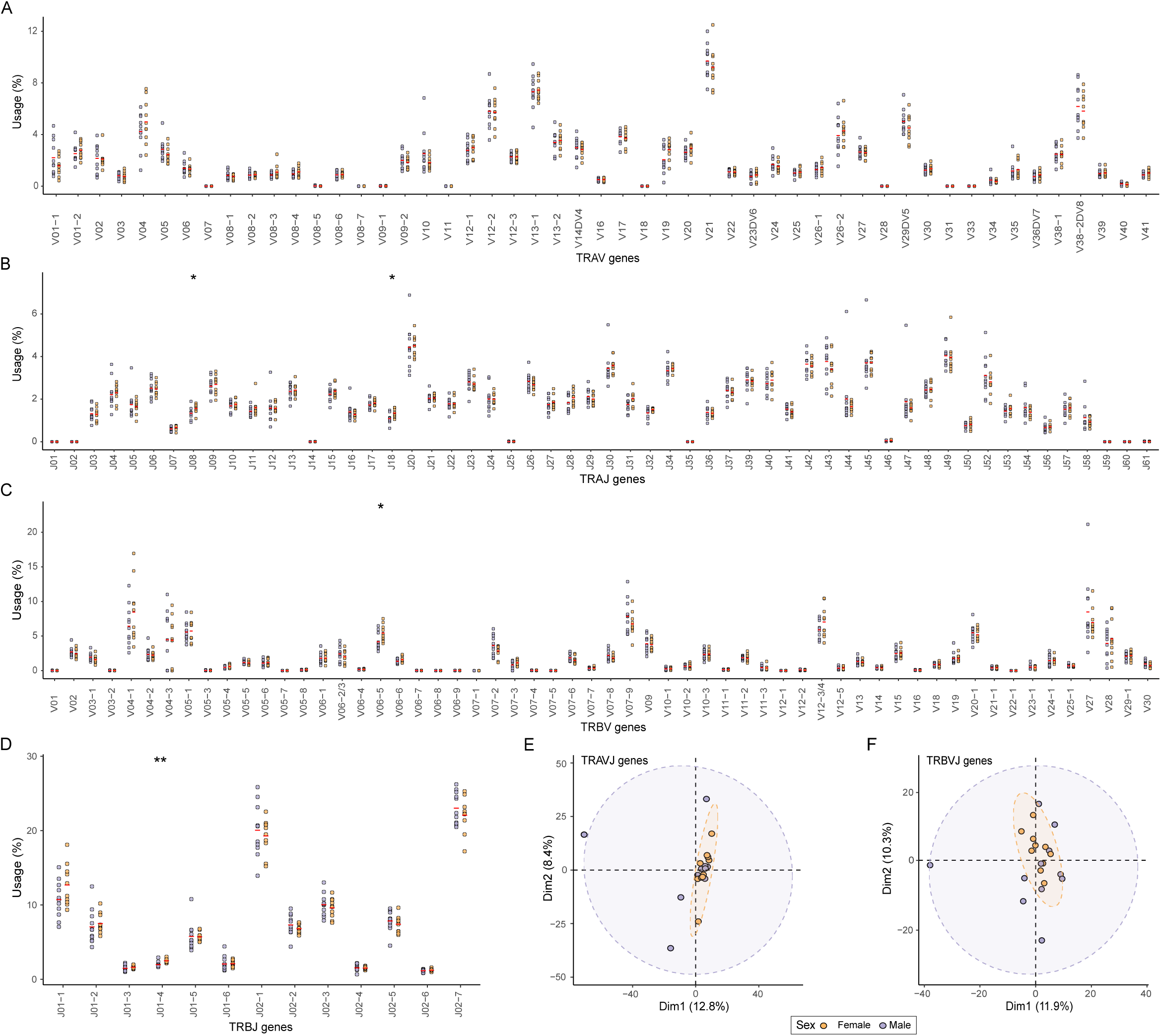
Some differential V and J gene usage in TCR repertoires of CD8 cells between males and females. (A-D) Frequencies of TRAV **(A)**, TRAJ **(B)**, TRBV **(C)**, and TRBJ **(D)** gene usage between males and females. Red lines represent the mean for each sex group. Statistical comparisons between groups were performed using the Wilcoxon test. Stars indicate statistical differences between males and females based on the p-value of the test (*: p < 0.05, **: p < 0.01). **(E-F)** Principal Component Analysis (PCA) derived from the distribution of the frequency usage of TRAV-TRAJ **(E)** and TRBV-TRBJ **(F)** gene associations across sex groups between males and females. Each point represents an individual. Ellipses show 95% confidence intervals. Males are depicted in violet and females in orange.

**Supplemental Figure 5:**
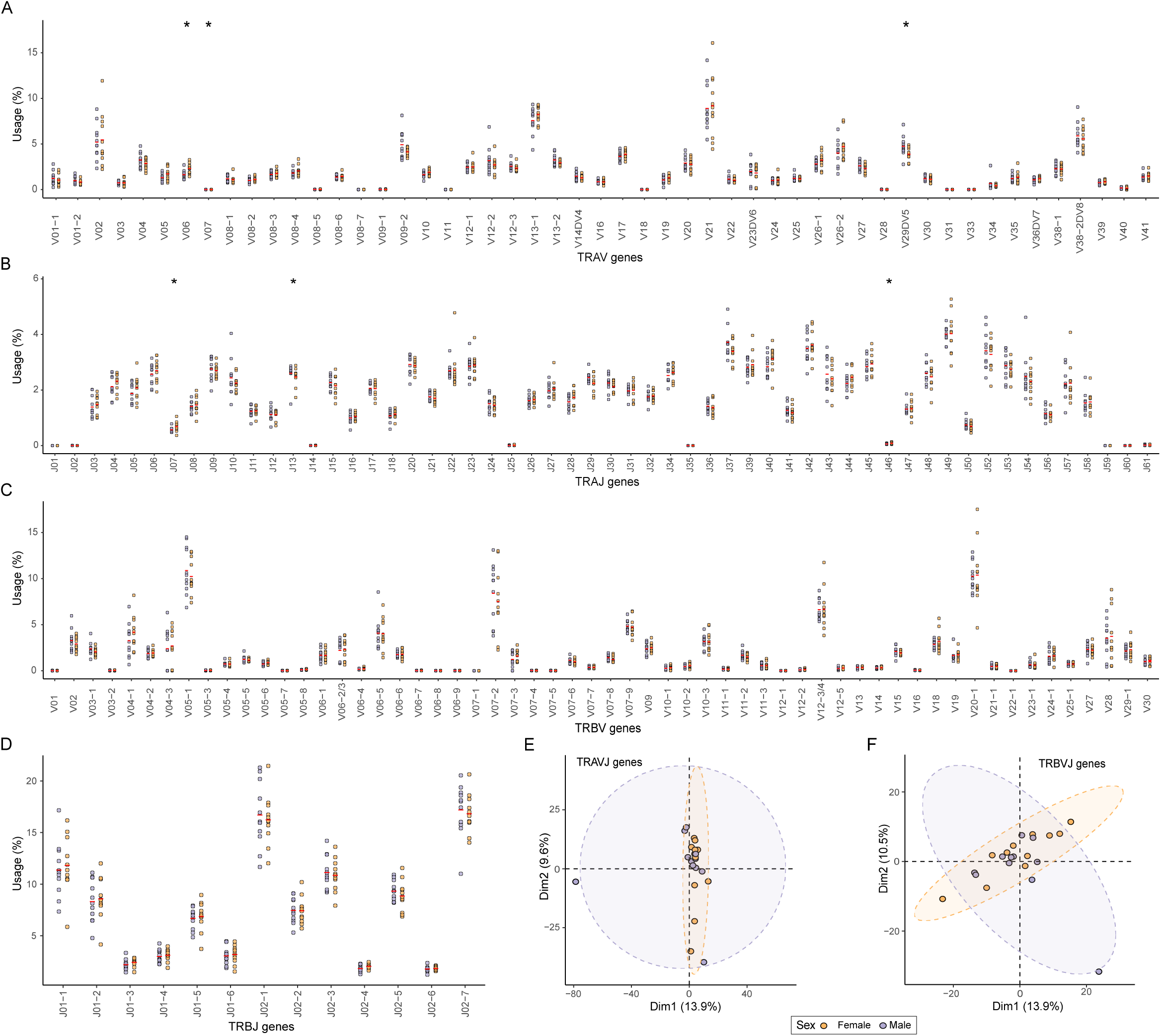
Some differential V and J gene usage in TCR repertoires of CD4 Teff cells between males and females. (A-D) Frequencies of TRAV **(A)**, TRAJ **(B)**, TRBV **(C)**, and TRBJ **(D)** gene usage between males and females. Red lines represent the mean for each sex group. Statistical comparisons between groups were performed using the Wilcoxon test. Stars indicate statistical differences between males and females based on the p-value of the test (*: p < 0.05, **: p < 0.01). **(E-F)** Principal Component Analysis (PCA) derived from the distribution of the frequency usage of TRAV-TRAJ **(E)** and TRBV-TRBJ **(F)** gene associations across sex groups between males and females. Each point represents an individual. Ellipses show 95% confidence intervals. Males are depicted in violet and females in orange.

**Supplemental Figure 6:**
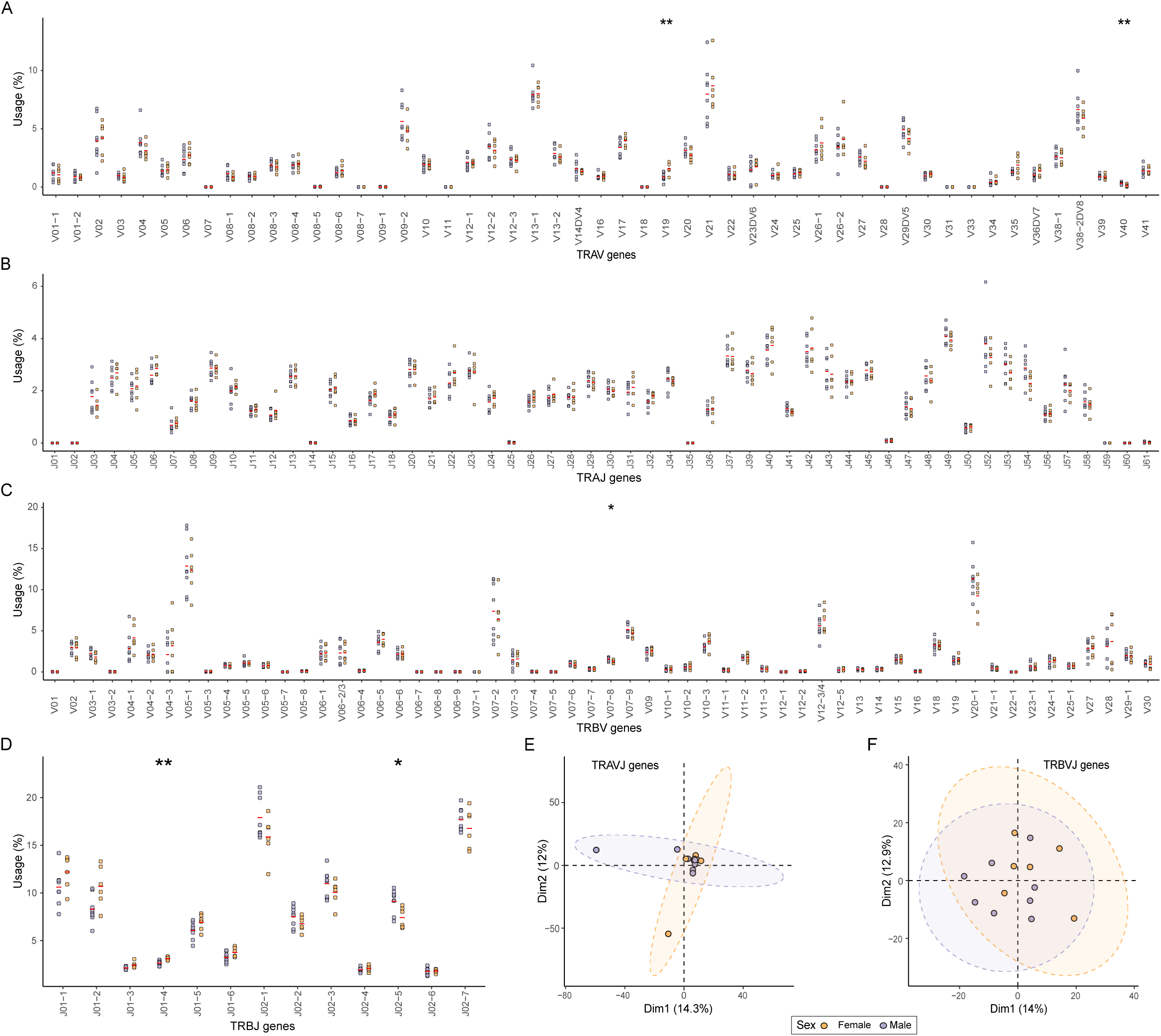
Some differential V and J gene usage in TCR repertoires of CD4 Treg cells between males and females. (A-D) Frequencies of TRAV **(A)**, TRAJ **(B)**, TRBV **(C)**, and TRBJ **(D)** gene usage between males and females. Red lines represent the mean for each sex group. Statistical comparisons between groups were performed using the Wilcoxon test. Stars indicate statistical differences between males and females based on the p-value of the test (*: p < 0.05, **: p < 0.01). **(E-F)** Principal Component Analysis (PCA) derived from the distribution of the usage frequency of TRAV-TRAJ **(E)** and TRBV-TRBJ **(F)** gene associations across sex groups between males and females. Each point represents an individual. Ellipses show 95% confidence intervals. Males are depicted in violet and females in orange.

**Supplemental Figure 7:**
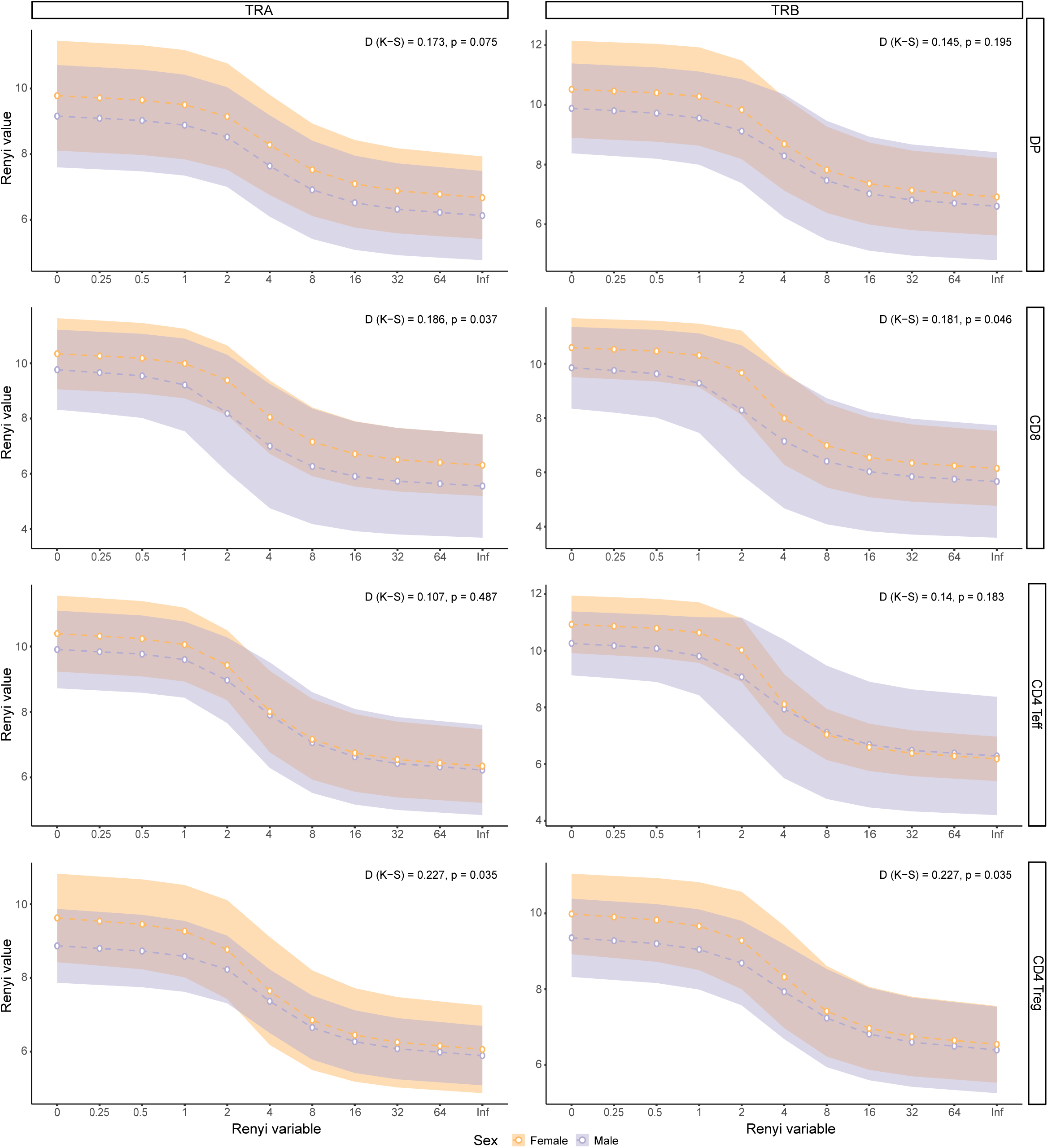
Minimal differences in diversity profile of CD8 and CD4 Treg thymic TCR repertoire between males and females. Diversity profile with Rényi diversity index values (α ranging from 0 to infinity) for all cell subsets, both TRA (left) and TRB (right). Clonotypes of each sample were rarefied fifty times to their effective diversity number [i.e. 𝑒^*shannon index*^] and the median of their Rényi values was used for analysis. Dotted lines and points show the mean value for each sex group, while the shaded area represents the standard deviation for each sex group. The overall shape of the curves was compared statistically using the Kolmogorov-Smirnov test, with the D value and associated p-value indicated.

**Supplemental Figure 8:**
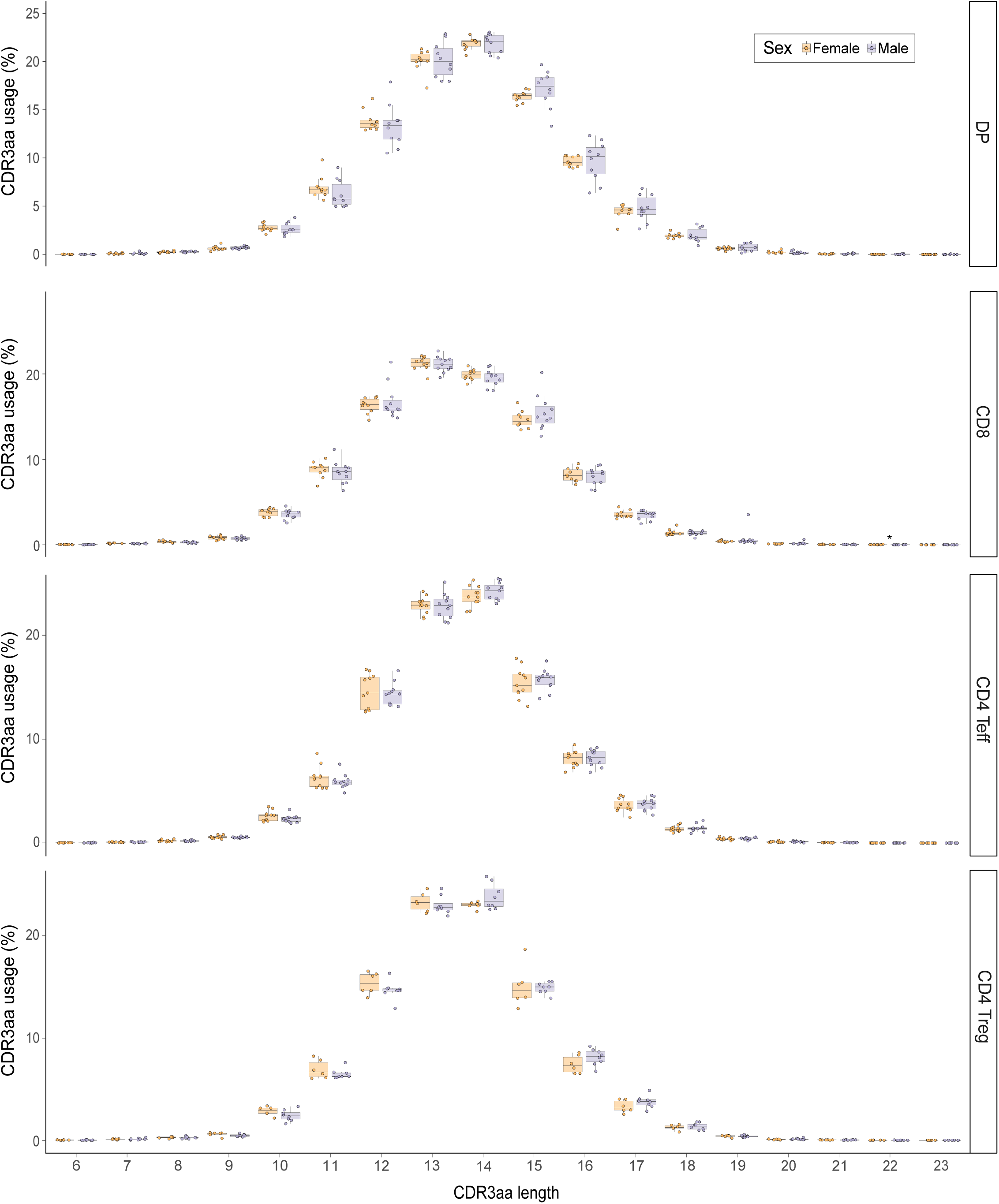
Comparable TRA CDR3aa length distribution between males and females. Distribution of CDR3aa length usage for TRA in all cell subsets. Stars indicate statistical differences between males and females based on the p-value of the Wilcoxon test (*: p < 0.05, **: p < 0.01). Males are depicted in violet and females in orange.

**Supplemental Figure 9:**
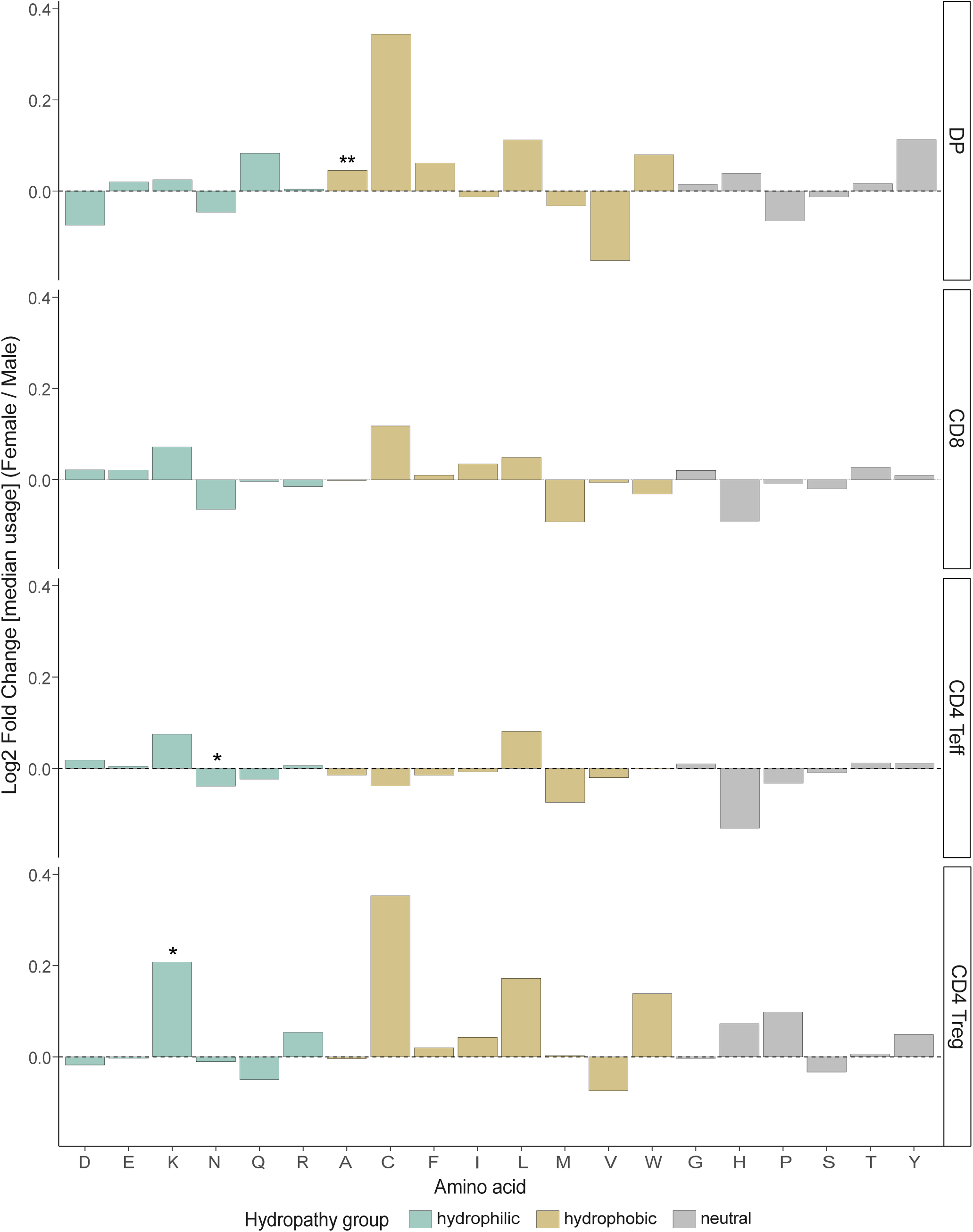
Some differences in TRA CDR3aa usage between males and females. Usage of each aa in the p108 to p114 CDR3 region represented as the log2 fold change of the median usage of females over males in TRA for each cell subset. A line at log2 fold change = 0 indicates the direction of the usage difference. Bars are colored according to the hydropathy class of the amino acid (neutral in gray, hydrophilic in blue-green and hydrophobic in gold). Stars indicate statistical differences between males and females based on the p-value of the Wilcoxon test (*: p < 0.05, **: p < 0.01).

**Supplemental Figure 10:**
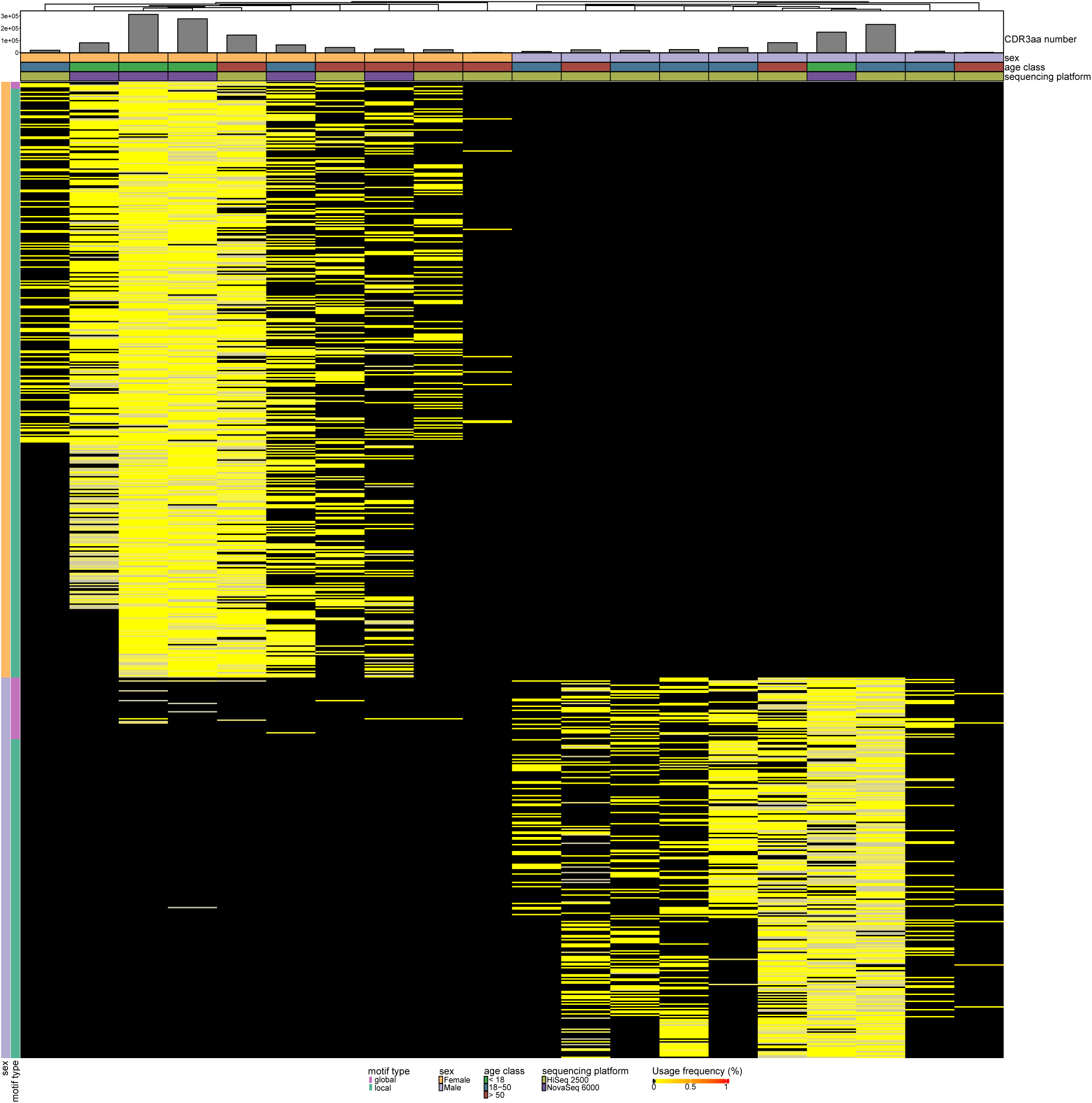
DP TRB thymic sex-associated motifs in our dataset. Different structural motifs in the CDR3 amino acid (CDR3aa) region found differentially expressed between males and females within our dataset. Local motifs refer to distinct amino acid sequences, whereas global motifs represent motif regions with a single variable amino acid position maintaining a BLOSUM62 score ≥ 0. The heatmap showcases the differential usage of TRB CDR3aa motifs between males and females in of DP cells. Hierarchical clustering reveals clear segregation of individuals by sex, with most CDR3aa motifs being almost exclusively expressed in one sex while absent in the other. Sample attributes are visualized as follows: (1) Sex: males in violet, females in orange; (2) Age class: children (<18 years) in green, young adults (18-50 years) in blue, older adults (>50 years) in red; (3) sequencing platform: HiSeq2500 samples in persimmon, NovaSeq 6000 samples in purple; (4) the total CDR3aa number. Motifs overexpressed in males are shown in violet, those overexpressed in females in orange; local motifs are indicated in blue-green and global motifs in magenta.

**Supplemental Figure 11:**
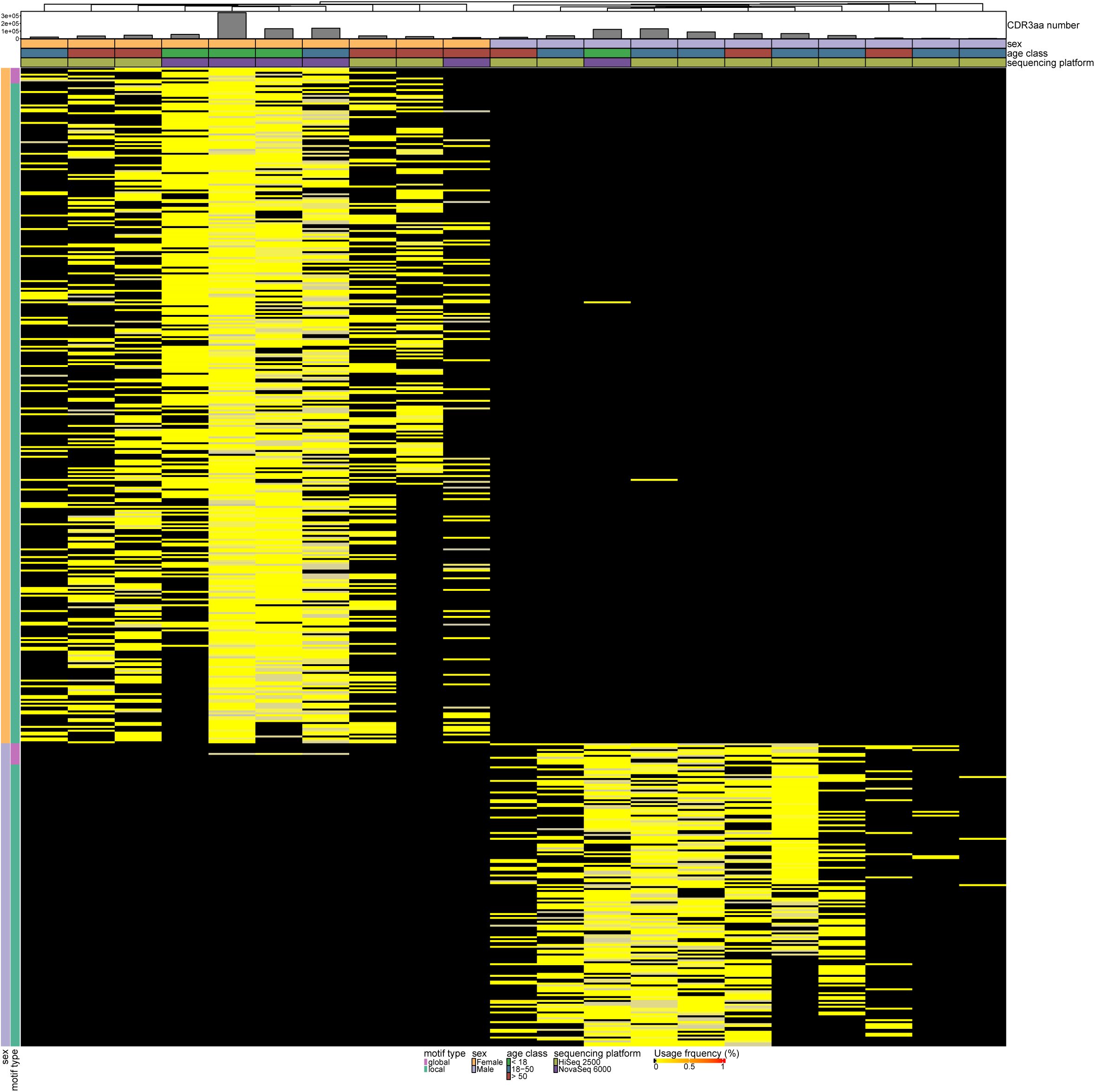
CD8 TRB thymic sex-associated motifs in our dataset. Different structural motifs in the CDR3 amino acid (CDR3aa) region found differentially expressed between males and females within our dataset. Local motifs refer to distinct amino acid sequences, whereas global motifs represent motif regions with a single variable amino acid position maintaining a BLOSUM62 score ≥ 0. The heatmap showcases the differential usage of TRB CDR3aa motifs between males and females in of CD8 cells. Hierarchical clustering reveals clear segregation of individuals by sex, with most CDR3aa motifs being almost exclusively expressed in one sex while absent in the other. Sample attributes are visualized as follows: (1) Sex: males in violet, females in orange; (2) Age class: children (<18 years) in green, young adults (18-50 years) in blue, older adults (>50 years) in red; (3) sequencing platform: HiSeq2500 samples in persimmon, NovaSeq 6000 samples in purple; (4) the total CDR3aa number. Motifs overexpressed in males are shown in violet, those overexpressed in females in orange; local motifs are indicated in blue-green and global motifs in magenta.

**Supplemental Figure 12:**
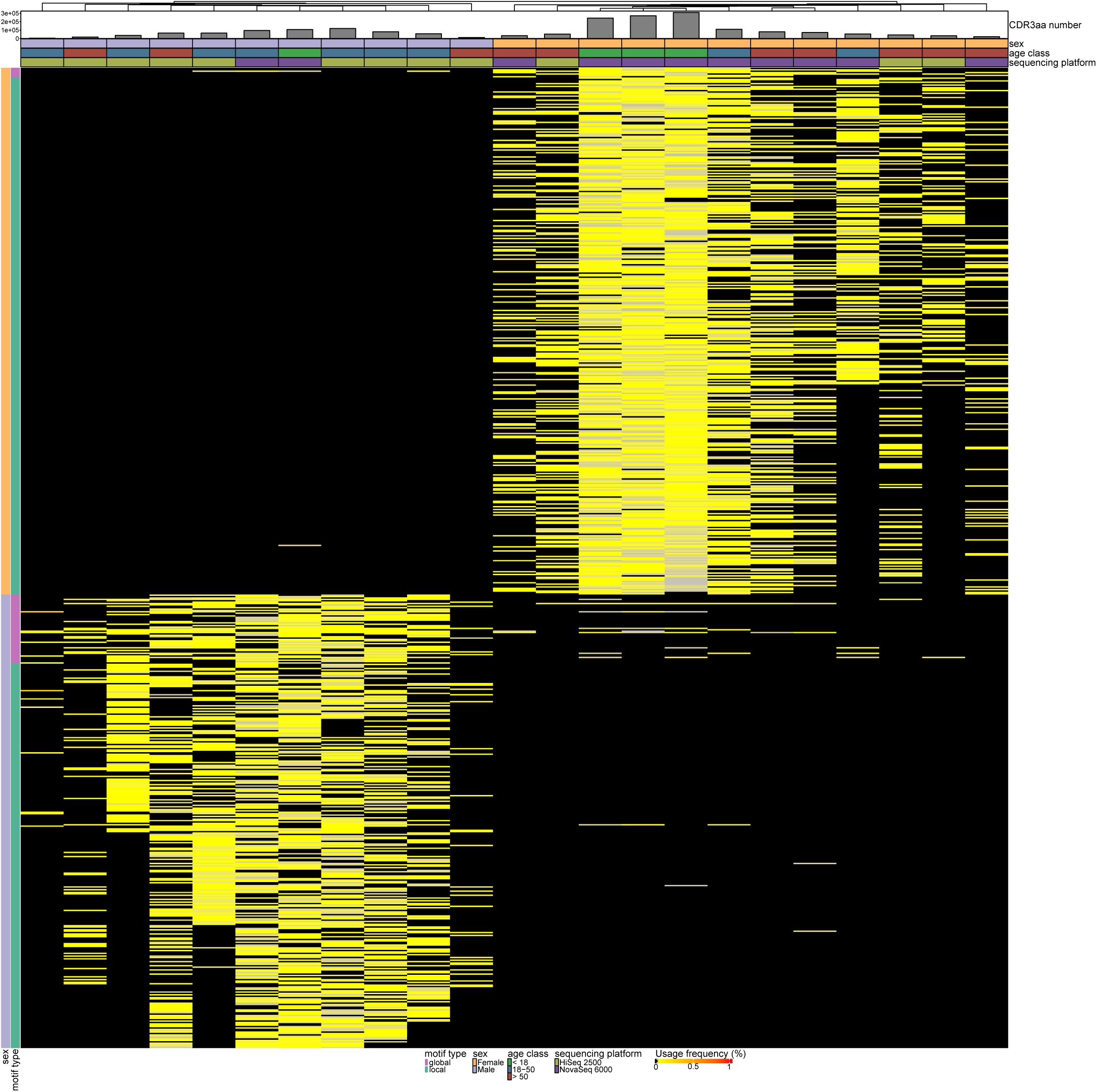
CD4 Teff TRB thymic sex-associated motifs in our dataset. Different structural motifs in the CDR3 amino acid (CDR3aa) region found differentially expressed between males and females within our dataset. Local motifs refer to distinct amino acid sequences, whereas global motifs represent motif regions with a single variable amino acid position maintaining a BLOSUM62 score ≥ 0. The heatmap showcases the differential usage of TRB CDR3aa motifs between males and females in of CD4 Teff cells. Hierarchical clustering reveals clear segregation of individuals by sex, with most CDR3aa motifs being almost exclusively expressed in one sex while absent in the other. Sample attributes are visualized as follows: (1) Sex: males in violet, females in orange; (2) Age class: children (<18 years) in green, young adults (18-50 years) in blue, older adults (>50 years) in red; (3) sequencing platform: HiSeq2500 samples in persimmon, NovaSeq 6000 samples in purple; (4) the total CDR3aa number. Motifs overexpressed in males are shown in violet, those overexpressed in females in orange; local motifs are indicated in blue-green and global motifs in magenta.

**Supplemental Figure 13:**
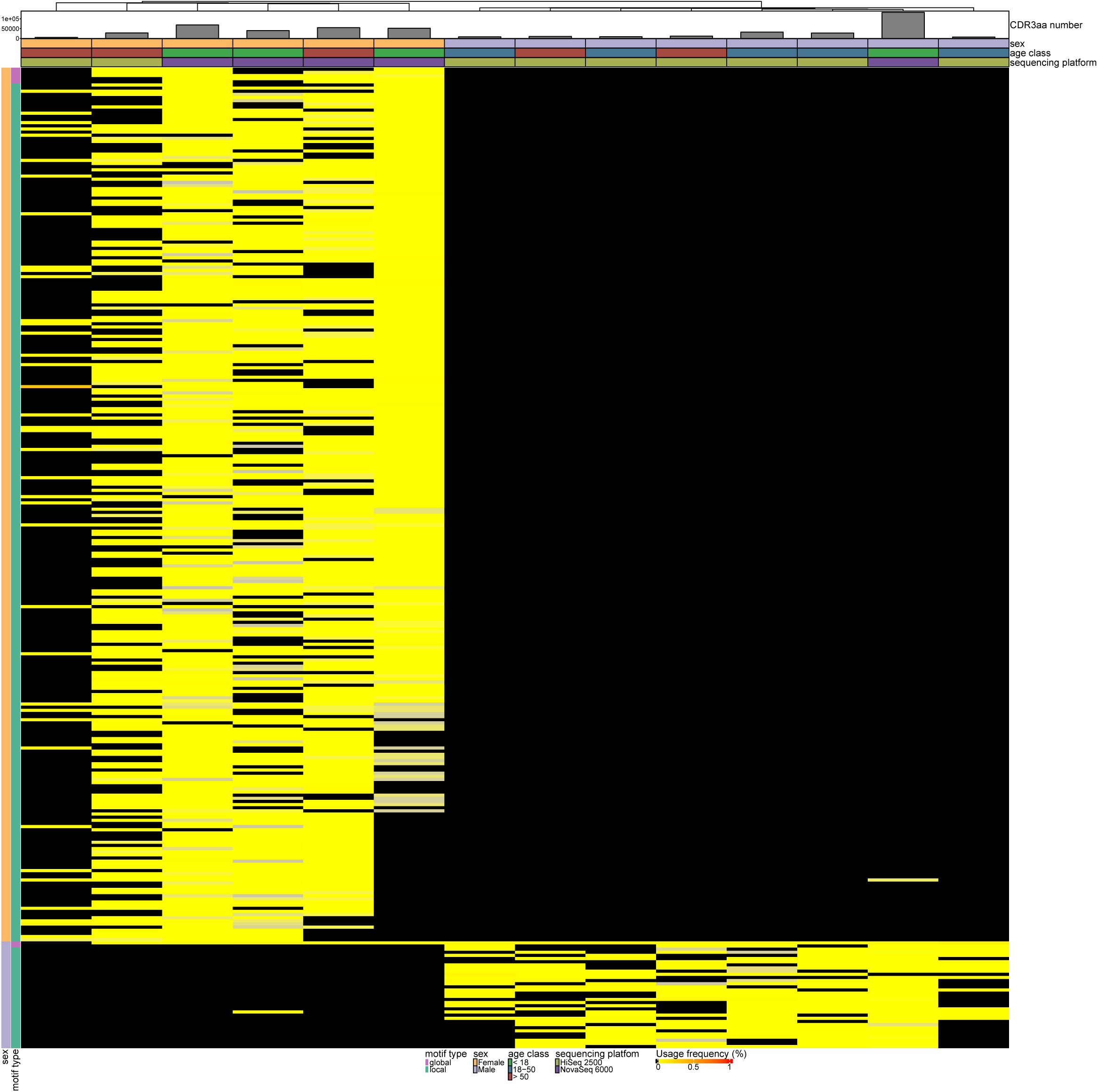
CD4 Treg TRB thymic sex-associated motifs in our dataset. Different structural motifs in the CDR3 amino acid (CDR3aa) region found differentially expressed between males and females within our dataset. Local motifs refer to distinct amino acid sequences, whereas global motifs represent motif regions with a single variable amino acid position maintaining a BLOSUM62 score ≥ 0. The heatmap showcases the differential usage of TRB CDR3aa motifs between males and females in of CD4 Treg cells. Hierarchical clustering reveals clear segregation of individuals by sex, with most CDR3aa motifs being almost exclusively expressed in one sex while absent in the other. Sample attributes are visualized as follows: (1) Sex: males in violet, females in orange; (2) Age class: children (<18 years) in green, young adults (18-50 years) in blue, older adults (>50 years) in red; (3) sequencing platform: HiSeq2500 samples in persimmon, NovaSeq 6000 samples in purple; (4) the total CDR3aa number. Motifs overexpressed in males are shown in violet, those overexpressed in females in orange; local motifs are indicated in blue-green and global motifs in magenta.

**Supplemental Figure 14:**
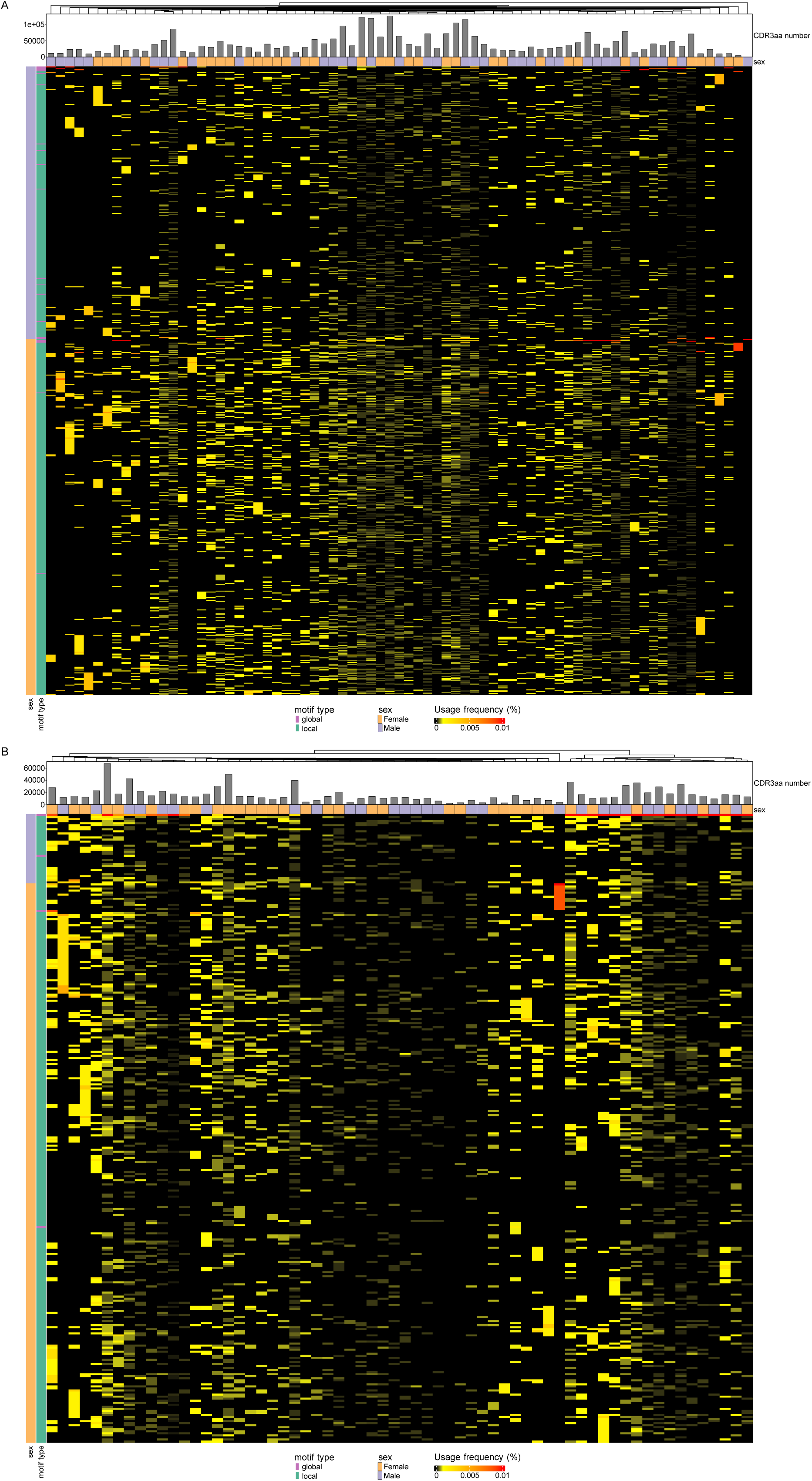
TRB thymic sex associated motifs are not differentially expressed between males and females in peripheral TCR repertoire. Heatmap representing TRB CDR3aa motif usage differentially expressed between males and females in CD4 Teff **(A)** and CD4 Treg **(B)** cells in a peripheral TCR repertoire dataset. Sex and total CDR3aa number are depicted by samples. Males are depicted in violet and females in orange and local motif in blue-green and global motifs in magenta.

**Supplemental Figure 15:**
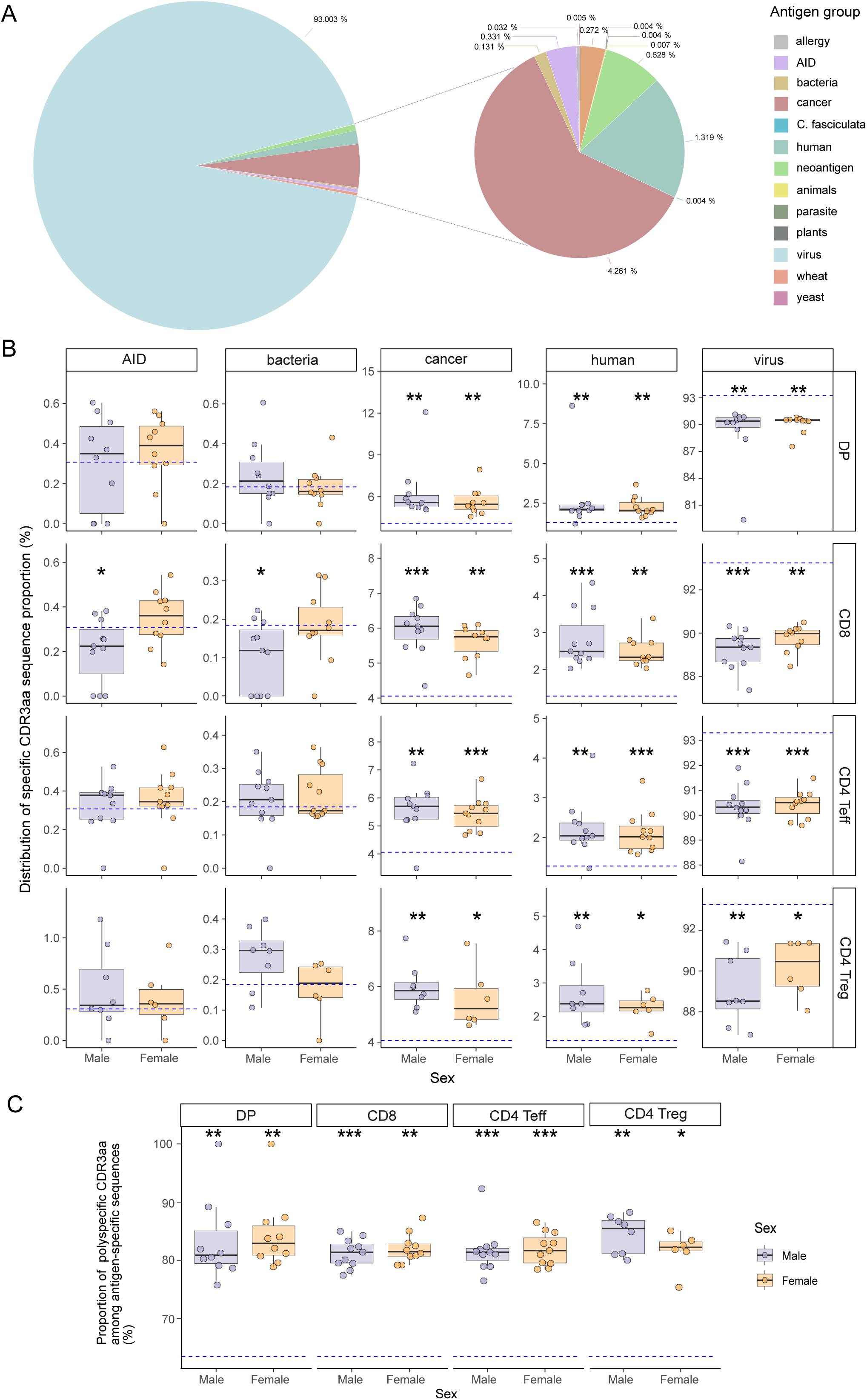
Enrichment of cancer associated, no-associated disease self-peptide specific and polyspecific TCRs, in our thymic dataset compared to initial curated database. **(A)** Representation of the specificity groups in the pooled and curated database. **(B)** Distribution of specific CDR3aa sequence proportions for each sample across autoimmune disease (AID), bacteria, cancer, human and virus groups compared to the reference database distributions for males and females. **(C)** Proportion of polyspecific CDR3aa among antigen-specific sequences of samples compared to the reference database for both sexes. Stars indicate statistical significance between groups (males or females) and the pooled, curated database based on Wilcoxon test p-values (*: p < 0.05, **: p < 0.01). Males are depicted in violet and females in orange.

**Supplemental Figure 16:**
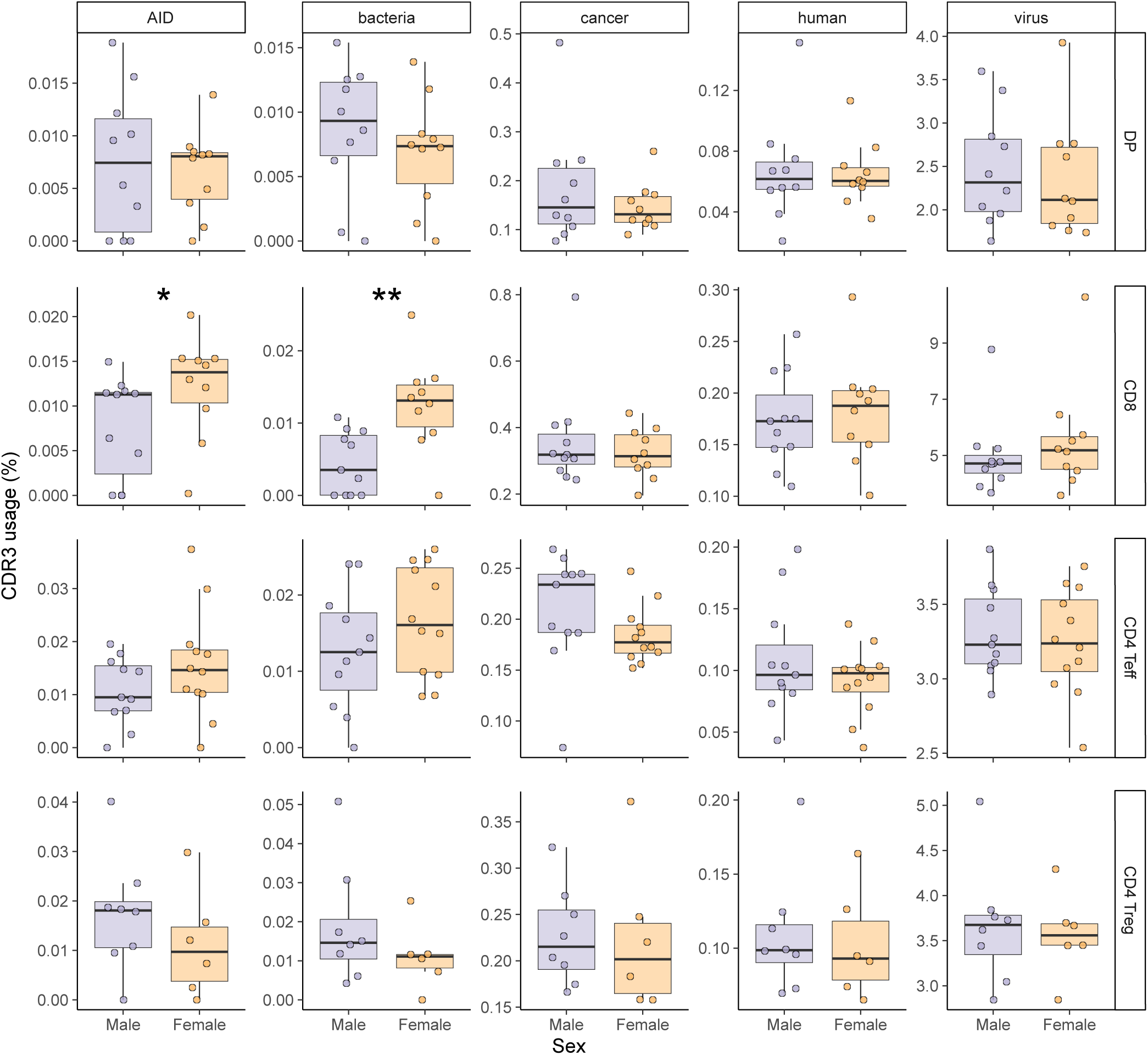
Enrichment of AID associated TCR and bacteria targeted TCR in CD8 female cells. CDR3aa usage of CDR3aa with a specificity in autoimmunity disease, bacteria, cancer, human and virus groups. Stars indicate statistical differences between males and females based on the p-value of the Wilcoxon test (*: p < 0.05, **: p < 0.01). Males are depicted in violet and females in orange.

**Supplemental Figure 17:**
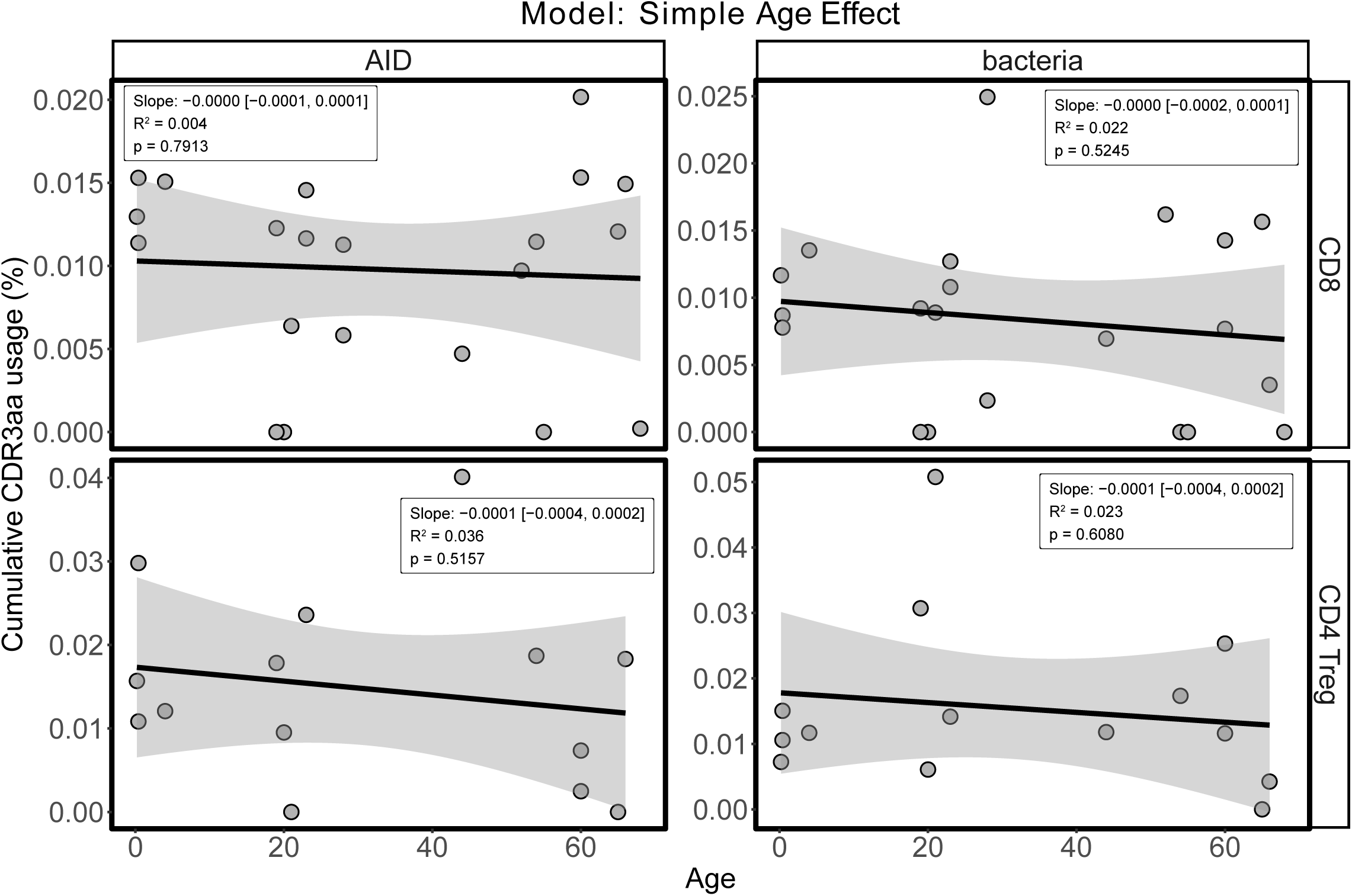
No association between age and thymic usage of CDR3 sequences specific to autoimmunity or bacterial antigens. Linear regression models assessing the effect of donor age on the cumulative usage (%) of TRB CDR3aa sequences with known specificity to self-antigens associated with autoimmunity (AID, left panels) or bacterial antigens (right panels), in CD8 SP (top) and CD4 Treg SP (bottom) thymic subsets. Each dot represents one donor. The shaded area indicates the 95% confidence interval around the regression line. Reported slope, 95% confidence interval, R², and p-values show no significant effect of age.

**Supplemental Table 1:** Minimal number of CDR3aa by cell subset.

|  | DP cells | CD8 cells | CD4 Teff | CD4 Treg |
| --- | --- | --- | --- | --- |
| TRA | 1013 | 2433 | 3314 | 2654 |
| TRB | 1429 | 3496 | 4551 | 5383 |

## References

1. Libert C, Dejager L, Pinheiro I. The X chromosome in immune functions: when a chromosome makes the difference. Nat Rev Immunol. 2010 Aug;10(8):594–604.

2. Golden LC, Voskuhl R. The Importance of Studying Sex Differences in Disease: The Example of Multiple Sclerosis. J Neurosci Res. 2017 Jan 2;95(1–2):633–43.

3. Sokka T, Toloza S, Cutolo M, Kautiainen H, Makinen H, Gogus F, et al. Women, men, and rheumatoid arthritis: analyses of disease activity, disease characteristics, and treatments in the QUEST-RA Study. Arthritis Res Ther. 2009;11(1):R7.

4. Li Y, Jerkic M, Slutsky AS, Zhang H. Molecular mechanisms of sex bias differences in COVID-19 mortality. Crit Care. 2020 July 9;24:405.

5. Hertz D, Schneider B. Sex differences in tuberculosis. Semin Immunopathol. 2019 Mar 1;41(2):225–37.

6. Klein SL, Morgan R. The impact of sex and gender on immunotherapy outcomes. Biol Sex Differ. 2020 May 4;11(1):24.

7. Lee HJ, Chang C. Recent advances in androgen receptor action. Cell Mol Life Sci CMLS. 2003 Aug 1;60(8):1613–22.

8. Chen P, Li B, Ou-Yang L. Role of estrogen receptors in health and disease. Front Endocrinol. 2022 Aug 18;13:839005.

9. Scarpin KM, Graham JD, Mote PA, Clarke CL. Progesterone action in human tissues: regulation by progesterone receptor (PR) isoform expression, nuclear positioning and coregulator expression. Nucl Recept Signal. 2009 Dec 31;7:e009.

10. Klein L, Kyewski B, Allen PM, Hogquist KA. Positive and negative selection of the T cell repertoire: what thymocytes see (and don’t see). Nat Rev Immunol. 2014 June;14(6):377–91.

11. Zhu ML, Bakhru P, Conley B, Nelson JS, Free M, Martin A, et al. Sex bias in CNS autoimmune disease mediated by androgen control of autoimmune regulator. Nat Commun. 2016 Apr 13;7(1):11350.

12. Dragin N, Bismuth J, Cizeron-Clairac G, Biferi MG, Berthault C, Serraf A, et al. Estrogen-mediated downregulation of AIRE influences sexual dimorphism in autoimmune diseases. J Clin Invest. 126(4):1525–37.

13. Stankiewicz LN, Salim K, Flaschner EA, Wang YX, Edgar JM, Durland LJ, et al. Sex-biased human thymic architecture guides T cell development through spatially defined niches. Dev Cell. 2024 Oct 3;S1534–5807(24)00539-2.

14. Murugan A, Mora T, Walczak AM, Callan CG. Statistical inference of the generation probability of T-cell receptors from sequence repertoires. Proc Natl Acad Sci. 2012 Oct 2;109(40):16161–6.

15. Yoshida K, Cologne JB, Cordova K, Misumi M, Yamaoka M, Kyoizumi S, et al. Aging-related changes in human T-cell repertoire over 20 years delineated by deep sequencing of peripheral T-cell receptors. Exp Gerontol. 2017 Oct 1;96:29–37.

16. Trofimov A, Brouillard P, Larouche JD, Séguin J, Laverdure JP, Brasey A, et al. Two types of human TCR differentially regulate reactivity to self and non-self antigens. iScience. 2022 Aug 17;25(9):104968.

17. Mika J, Yoshida K, Kusunoki Y, Candéias SM, Polanska J. Sex-and age-specific aspects of human peripheral T-cell dynamics. Front Immunol. 2023 Oct 13;14:1224304.

18. Gong M, Li X, Zheng A, Xu H, Xie S, Yan R, et al. Age-related changes in the TRB and IGH repertoires in healthy adult males and females. Immunol Lett. 2021 Dec 1;240:71–6.

19. Qi Q, Liu Y, Cheng Y, Glanville J, Zhang D, Lee JY, et al. Diversity and clonal selection in the human T-cell repertoire. Proc Natl Acad Sci U S A. 2014 Sept 9;111(36):13139–44.

20. Davis MM, Bjorkman PJ. T-cell antigen receptor genes and T-cell recognition. Nature. 1988 Aug;334(6181):395–402.

21. Wang F, Ono T, Kalergis AM, Zhang W, DiLorenzo TP, Lim K, et al. On defining the rules for interactions between the T cell receptor and its ligand: a critical role for a specific amino acid residue of the T cell receptor beta chain. Proc Natl Acad Sci U S A. 1998 Apr 28;95(9):5217–22.

22. Yu K, Shi J, Lu D, Yang Q. Comparative analysis of CDR3 regions in paired human αβ CD8 T cells. FEBS Open Bio. 2019 July 12;9(8):1450–9.

23. Carter JA, Preall JB, Grigaityte K, Goldfless SJ, Jeffery E, Briggs AW, et al. Single T Cell Sequencing Demonstrates the Functional Role of αβ TCR Pairing in Cell Lineage and Antigen Specificity. Front Immunol. 2019;10:1516.

24. Garcia KC, Degano M, Pease LR, Huang M, Peterson PA, Teyton L, et al. Structural basis of plasticity in T cell receptor recognition of a self peptide-MHC antigen. Science. 1998 Feb 20;279(5354):1166–72.

25. Pommié C, Levadoux S, Sabatier R, Lefranc G, Lefranc MP. IMGT standardized criteria for statistical analysis of immunoglobulin V-REGION amino acid properties. J Mol Recognit. 2004;17(1):17–32.

26. Khosravi-Maharlooei M, Obradovic A, Misra A, Motwani K, Holzl M, Seay HR, et al. Crossreactive public TCR sequences undergo positive selection in the human thymic repertoire. J Clin Invest. 2019 Mar 28;129(6):2446–62.

27. Lu J, Van Laethem F, Bhattacharya A, Craveiro M, Saba I, Chu J, et al. Molecular constraints on CDR3 for thymic selection of MHC-restricted TCRs from a random pre-selection repertoire. Nat Commun. 2019 Mar 4;10:1019.

28. Stadinski BD, Shekhar K, Gómez-Touriño I, Jung J, Sasaki K, Sewell AK, et al. Hydrophobic CDR3 residues promote the development of self-reactive T cells. Nat Immunol. 2016 Aug;17(8):946–55.

29. Elhanati Y, Murugan A, Callan CG, Mora T, Walczak AM. Quantifying selection in immune receptor repertoires. Proc Natl Acad Sci U S A. 2014 July 8;111(27):9875–80.

30. Meysman P, De Neuter N, Gielis S, Bui Thi D, Ogunjimi B, Laukens K. On the viability of unsupervised T-cell receptor sequence clustering for epitope preference. Bioinformatics. 2019 May 1;35(9):1461–8.

31. Springer I, Tickotsky N, Louzoun Y. Contribution of T Cell Receptor Alpha and Beta CDR3, MHC Typing, V and J Genes to Peptide Binding Prediction. Front Immunol. 2021 Apr 26;12:664514.

32. Korpela D, Jokinen E, Dumitrescu A, Huuhtanen J, Mustjoki S, Lähdesmäki H. EPIC-TRACE: predicting TCR binding to unseen epitopes using attention and contextualized embeddings. Bioinformatics. 2023 Dec 9;39(12):btad743.

33. Huang H, Wang C, Rubelt F, Scriba TJ, Davis MM. Analyzing the Mycobacterium tuberculosis immune response by T-cell receptor clustering with GLIPH2 and genome-wide antigen screening. Nat Biotechnol. 2020 Oct;38(10):1194–202.

34. Henikoff S, Henikoff JG. Amino acid substitution matrices from protein blocks. Proc Natl Acad Sci U S A. 1992 Nov 15;89(22):10915–9.

35. Heikkilä N, Kleino I, Vanhanen R, Yohannes DA, Mattila IP, Saramäki J, et al. Characterization of human T cell receptor repertoire data in eight thymus samples and four related blood samples. Data Brief. 2021 Apr 1;35:106751.

36. Mattila J, Sormunen S, Heikkilä N, Mattila IP, Saramäki J, Arstila TP. Analysis of thymic generation of shared T-cell receptor α repertoire associated with recognition of tumor antigens shows no preference for neoantigens over wild-type antigens. Cancer Med. 2023 June;12(12):13486–96.

37. Vita R, Mahajan S, Overton JA, Dhanda SK, Martini S, Cantrell JR, et al. The Immune Epitope Database (IEDB): 2018 update. Nucleic Acids Res. 2018 Oct 24;47(Database issue):D339.

38. Tickotsky N, Sagiv T, Prilusky J, Shifrut E, Friedman N. McPAS-TCR: a manually curated catalogue of pathology-associated T cell receptor sequences. Bioinforma Oxf Engl. 2017 Sept 15;33(18):2924–9.

39. Shugay M, Bagaev DV, Zvyagin IV, Vroomans RM, Crawford JC, Dolton G, et al. VDJdb: a curated database of T-cell receptor sequences with known antigen specificity. Nucleic Acids Res. 2018 Jan 4;46(D1):D419–27.

40. Rosati E, Dowds CM, Liaskou E, Henriksen EKK, Karlsen TH, Franke A. Overview of methodologies for T-cell receptor repertoire analysis. BMC Biotechnol. 2017 July 10;17(1):61.

41. Wooldridge L, Ekeruche-Makinde J, van den Berg HA, Skowera A, Miles JJ, Tan MP, et al. A single autoimmune T cell receptor recognizes more than a million different peptides. J Biol Chem. 2012 Jan 6;287(2):1168–77.

42. Hadrup SR, Bakker AH, Shu CJ, Andersen RS, van Veluw J, Hombrink P, et al. Parallel detection of antigen-specific T-cell responses by multidimensional encoding of MHC multimers. Nat Methods. 2009 July;6(7):520–6.

43. Quiniou V, Barennes P, Mhanna V, Stys P, Vantomme H, Zhou Z, et al. Human thymopoiesis produces polyspecific CD8+ α/β T cells responding to multiple viral antigens. eLife. 2023 Mar 30;12:e81274.

44. van de Sandt CE, Nguyen THO, Gherardin NA, Crawford JC, Samir J, Minervina AA, et al. Newborn and child-like molecular signatures in older adults stem from TCR shifts across human lifespan. Nat Immunol. 2023 Nov;24(11):1890–907.

45. Schneider-Hohendorf T, Görlich D, Savola P, Kelkka T, Mustjoki S, Gross CC, et al. Sex bias in MHC I-associated shaping of the adaptive immune system. Proc Natl Acad Sci U S A. 2018 Feb 27;115(9):2168–73.

46. Roca AM, Chobrutskiy BI, Callahan BM, Blanck G. T-cell receptor V and J usage paired with specific HLA alleles associates with distinct cervical cancer survival rates. Hum Immunol. 2019 Apr;80(4):237–42.

47. Hou X, Wei W, Zhang J, Liu Z, Wang G, Yang X, et al. Characterisation of T and B cell receptor repertoire in patients with systemic lupus erythematosus. Clin Exp Rheumatol. 2023 Nov;41(11):2216–23.

48. Rojas M, Restrepo-Jiménez P, Monsalve DM, Pacheco Y, Acosta-Ampudia Y, Ramírez-Santana C, et al. Molecular mimicry and autoimmunity. J Autoimmun. 2018 Dec 1;95:100–23.

49. Repac J, Božić B, Božić Nedeljković B. Microbes as triggers and boosters of Type 1 Diabetes - Mediation by molecular mimicry. Diabetes Res Clin Pract. 2023 Aug;202:110824.

50. Girdhar K, Huang Q, Chow IT, Vatanen T, Brady C, Raisingani A, et al. A gut microbial peptide and molecular mimicry in the pathogenesis of type 1 diabetes. Proc Natl Acad Sci U S A. 2022 Aug 2;119(31):e2120028119.

51. Petersen J, Ciacchi L, Tran MT, Loh KL, Kooy-Winkelaar Y, Croft NP, et al. T cell receptor cross-reactivity between gliadin and bacterial peptides in celiac disease. Nat Struct Mol Biol. 2020 Jan;27(1):49–61.

52. Vazquez DS, Schilbert HM, Dodero VI. Molecular and Structural Parallels between Gluten Pathogenic Peptides and Bacterial-Derived Proteins by Bioinformatics Analysis. Int J Mol Sci. 2021 Aug 27;22(17):9278.

53. Tai N, Peng J, Liu F, Gulden E, Hu Y, Zhang X, et al. Microbial antigen mimics activate diabetogenic CD8 T cells in NOD mice. J Exp Med. 2016 Sept 19;213(10):2129–46.

54. Hebbandi Nanjundappa R, Sokke Umeshappa C, Geuking MB. The impact of the gut microbiota on T cell ontogeny in the thymus. Cell Mol Life Sci. 2022 Apr;79(4):221.

55. Markle JGM, Frank DN, Mortin-Toth S, Robertson CE, Feazel LM, Rolle-Kampczyk U, et al. Sex differences in the gut microbiome drive hormone-dependent regulation of autoimmunity. Science. 2013 Mar 1;339(6123):1084–8.

56. Ghoreyshi Z, George J. Quantitative approaches for decoding the specificity of the human T cell repertoire. Front Immunol. 2023 Sept 7;14.

57. Barennes P, Quiniou V, Shugay M, Egorov ES, Davydov AN, Chudakov DM, et al. Benchmarking of T cell receptor repertoire profiling methods reveals large systematic biases. Nat Biotechnol. 2021 Feb;39(2):236–45.

58. Bolotin DA, Poslavsky S, Mitrophanov I, Shugay M, Mamedov IZ, Putintseva EV, et al. MiXCR: software for comprehensive adaptive immunity profiling. Nat Methods. 2015 May;12(5):380–1.

59. Omer A, Peres A, Rodriguez OL, Watson CT, Lees W, Polak P, et al. T cell receptor beta germline variability is revealed by inference from repertoire data. Genome Med. 2022 Jan 7;14(1):2.

60. Chaara W, Gonzalez-Tort A, Florez LM, Klatzmann D, Mariotti-Ferrandiz E, Six A. RepSeq Data Representativeness and Robustness Assessment by Shannon Entropy. Front Immunol. 2018 May 15;9:1038.

61. Shannon CE. The mathematical theory of communication. 1963. MD Comput Comput Med Pract. 1997;14(4):306–17.

62. Rényi A. On Measures of Entropy and Information. In: Proceedings of the Fourth Berkeley Symposium on Mathematical Statistics and Probability, Volume 1: Contributions to the Theory of Statistics. University of California Press; 1961. p. 547–62.

63. Lefranc MP, Pommié C, Ruiz M, Giudicelli V, Foulquier E, Truong L, et al. IMGT unique numbering for immunoglobulin and T cell receptor variable domains and Ig superfamily V-like domains. Dev Comp Immunol. 2003 Jan;27(1):55–77.

64. Kyte J, Doolittle RF. A simple method for displaying the hydropathic character of a protein. J Mol Biol. 1982 May 5;157(1):105–32.

65. Marcou Q, Mora T, Walczak AM. High-throughput immune repertoire analysis with IGoR. Nat Commun. 2018 Feb 8;9(1):561.

66. Sethna Z, Elhanati Y, Callan CG, Walczak AM, Mora T. OLGA: fast computation of generation probabilities of B-and T-cell receptor amino acid sequences and motifs. Bioinforma Oxf Engl. 2019 Sept 1;35(17):2974–81.

67. Madi A, Poran A, Shifrut E, Reich-Zeliger S, Greenstein E, Zaretsky I, et al. T cell receptor repertoires of mice and humans are clustered in similarity networks around conserved public CDR3 sequences. eLife. 2017 July 21;6:e22057.

68. Lorenzon R, Mariotti-Ferrandiz E, Aheng C, Ribet C, Toumi F, Pitoiset F, et al. Clinical and multi-omics cross-phenotyping of patients with autoimmune and autoinflammatory diseases: the observational TRANSIMMUNOM protocol. BMJ Open. 2018 Aug 30;8(8):e021037.

69. Vita R, Mahajan S, Overton JA, Dhanda SK, Martini S, Cantrell JR, et al. The Immune Epitope Database (IEDB): 2018 update. Nucleic Acids Res. 2019 Jan 8;47(D1):D339–43.

70. Jouannet C, Vantomme H, Gouge KL, Klatzmann D, Mariotti-Ferrandiz E. Benchmarking unsupervised methods for inferring TCR specificity. NAR Genomics Bioinforma. 2025 Dec 1;7(4):lqaf150.

71. Yi X, Liao Y, Wen B, Li K, Dou Y, Savage SR, et al. caAtlas: An immunopeptidome atlas of human cancer. iScience. 2021 Oct 22;24(10):103107.

